# Schwann Cell–Specific TDP-43 Rescue Improves Peripheral Nerve Myelin Pathology Without Altering Motor Behaviour in a Mouse Model of ALS

**DOI:** 10.64898/2026.07.20.739467

**Authors:** Katherine N. Lewis, Aaron Andres, Georgina Craig, Andrew P. Tosolini, Robin McAllen, Shyuan T. Ngo, David G. Gonsalvez, Bradley J. Turner, Samantha K. Barton

## Abstract

Amyotrophic lateral sclerosis (ALS) is a terminal disease caused by motor neuron loss. Schwann cells, the myelinating cells of the peripheral nervous system, metabolically and structurally support neurons. ALS patients exhibit Schwann cell pathology, such as TDP-43 proteinopathy, therefore Schwann cell dysfunction may contribute to disease progression. Here, we have characterised myelinating Schwann cell pathology in a TDP-43^Q331K^ (TDP-43) transgenic mouse model of ALS. We also crossed the floxxed TDP-43 mouse with a myelin protein zero (P0)-cre mouse to excise the transgene from Schwann cells alone (P0-cre/TDP-43) to assess rescue. Compared to wild-type (WT) littermates, 10 mo TDP-43 mice exhibited changes to myelin architecture, including loss of myelin binding proteins at the paranodes, decreased node of Ranvier length, and non-compact, degenerating myelin. In P0-cre/TDP-43 mice these myelin disruptions were rescued. However, this improved histology did not lead to a functional rescue, with both P0-cre/TDP-43 and TDP-43 mice exhibiting slowed sciatic nerve conduction and worsened motor behaviour. Further histological analyses revealed that Büngner Schwann cells, a subtype of Schwann cells triggered by neuronal injury, were activated in both TDP-43 and P0-cre/TDP-43 mice. Activation of Büngner Schwann cells can trigger damaging inflammation through the recruitment of macrophages, which can hinder motor and electrophysiological performance, potentially underpinning the lack of functional rescue in the P0-cre/TDP-43. We established that the rescue of Schwann cells indeed protects myelin in this ALS model, however understanding how Büngner Schwann cells exacerbate neuronal pathology is essential for developing effective therapeutics that can improve functional output.

**Significance Statement:** Amyotrophic lateral sclerosis (ALS) is a fatal neurodegenerative disease with no cure and limited treatments. Given that the average patient life expectancy is 3-5 years following diagnosis, finding novel treatment targets is of the utmost importance. Recent research has revealed that non-neuronal cells, such as Schwann cells, contribute to the disease, however the extent of their pathology remains elusive. Investigating Schwann cell and peripheral myelin pathology in ALS may lead to the identification of previously unrecognized disease mechanisms, opening novel avenues for therapeutic development. Identifying approaches through which to target glial and neuronal pathology concurrently would enable more holistic treatment of the various aspects of ALS pathobiology to improve patient outcomes.

## Introduction

Amyotrophic lateral sclerosis (ALS) is a fatal neurodegenerative disease characterised by the death of upper motor neurons within the central nervous system (CNS) and lower motor neurons within the peripheral nervous system (PNS) (Al-Chalabi et al., 2016). Common to 97% of all ALS patients are TDP-43 proteinopathies (Ling et al., 2015; Ling et al., 2013) whereby the essential DNA and RNA binding protein, TDP-43, encoded by the *TARDBP* gene (Cheng et al., 2024; Dong & Chen, 2018), becomes aberrantly regulated, causing cell death via both loss and gain of function mechanisms (Balendra et al., 2025; Dong & Chen, 2018). TDP-43 pathology is not unique to degenerating motor neurons and pathological TDP-43 aggregates have also been identified in other cell types including Schwann cells in ALS patients (Nakamura-Shindo et al., 2020; Riva et al., 2022). Schwann cells are the myelin producing cells of the PNS that function to insulate peripheral axons in a lipid-dense sheath called myelin (Nave & Werner, 2021), allowing for efficient nerve conduction along the axon (Belin et al., 2017; Uncini et al., 2024), and provide essential metabolic and trophic support to the axon (Bouçanova et al., 2021; Deck et al., 2022; Taveggia & Feltri, 2022). Schwann cells also play integral roles in facilitating axon repair after injury (Jessen & Mirsky, 2016), where they reprogram into ‘repair’ or Büngner Schwann cells, rapidly proliferate (Gomez-Sanchez et al., 2017), assist in clearance of debris (Gomez-Sanchez et al., 2015), recruit immune cells and modulate cytokines (Berner et al., 2022; Dubový et al., 2014; Martini et al., 2008), ultimately allowing the formation of new axonal tracts and axon regeneration (Jessen & Mirsky, 2016). However, in instances of chronic activation, Büngner Schwann cells can cause detrimental inflammatory effects (Berner et al., 2022; Gomez-Sanchez et al., 2015; Hartlehnert et al., 2017; Trias et al., 2020; Wagner & Myers, 1996). Despite the critical roles of Schwann cells in maintaining lower motor neuron health, the intrinsic contributions of these cells in ALS remains elusive.

To date, studies investigating the roles of Schwann cells in ALS have largely focused on mouse models carrying *superoxide dismutase 1* (*SOD1*) mutations yet have yielded conflicting results. In SOD1^G85R^ mice where the human transgene was excised from Schwann cells alone (using a P0 promoter), P0-cre/SOD1^G85R^ mice experienced delayed disease onset and extended survival (Wang et al., 2012). In contrast, when the study was repeated in SOD1^G37R^ mice, P0-cre/SOD1^G37R^ mice had accelerated late-stage disease progression (Lobsiger et al., 2009). These opposing results highlight how little is understood regarding the contributions of myelinating Schwann cells to ALS pathology. Second to this, these data are not entirely reflective of TDP-43-relevant ALS due to the disparate mechanisms of SOD1 pathology in disease. In a context of TDP-43, selective deletion of the endogenous *TARDBP* gene in Schwann cells in otherwise healthy mice resulted in a loss of myelin binding proteins at the paranode, directly causing defects in motor behaviour and reduced nerve conduction velocity (Chang et al., 2021). This suggests that dysfunctional Schwann cell encoded TDP-43 can directly impact motor performance and electrophysiological properties. While conditional deletion of *TARDBP* provides integral knowledge regarding the relationship between TDP-43 biology and Schwann cells, complete deletion of *TARDBP* does not accurately reflect the missense mutation-linked disease observed in human ALS. Thus, the use of alternative models is needed to characterise the relationship between TDP-43 pathology, Schwann cells, and ALS.

Here, we aimed to characterise the contributions of Schwann cells to ALS disease phenotypes using a TDP-43^Q331K^ transgenic mouse model of ALS, which causes the overexpression of mutant TDP-43 protein in neurons and glia (Arnold et al., 2013). We crossed the floxxed TDPIZI43^Q331K^ transgenic mice with myelin protein zero (P0)-cre mice (Feltri et al., 1999), resulting in Schwann cell-specific excision of the TDPIZI43 transgene, and assessed the intrinsic involvement of Schwann cells in this ALS mouse.

## Results

### TDP-43 Mice Exhibit Altered PNS Nodal and Paranodal Pathology that is Rescued in P0-cre/TDP-43 Mice

Since the heterozygous TDP-43^Q331K^ mouse has loxP sites enabling transgene excision, we crossed these mice with the P0-cre mouse strain (Feltri et al., 1999) which has also been used previously in SOD1 related ALS studies (Lobsiger et al., 2009; Wang et al., 2012), and is well established to have Schwann cell specific knock-down (Feltri et al., 1999). To ensure peripheral knock-down of TDP-43 protein in our cohort, Western blotting showed a 45.82 % ± 3.78 reduction in total TDP-43 protein in P0-cre/TDP-43 sciatic nerves, compared to TDP-43 sciatic nerves, at 2 mo, which aligns with previous findings (Lobsiger et al., 2009) (Supplementary Fig. 1a). This knock-down was sustained in 10 mo P0-cre/TDP-43 mice (Supplementary Fig. 1b).

Next, we aimed to characterise myelinating Schwann cell pathology in 10 mo TDP-43 mice and whether it was rescued in P0-cre/TDP-43 mice. To ascertain paranodal morphology, we stained for the paranodal marker Caspr, which in healthy contexts bilaterally flanks both sides of the node of Ranvier (which we immunolabelled with Nfasc; Fig. 1A). Surprisingly, we saw several instances where Nfasc^+^ nodes were present without Caspr^+^ paranodes (termed ‘missing’; Fig. 1B). When compared to WT sciatic nerves, TDP-43 sciatic nerves had an approximate two-fold increase in the number of Nfasc^+^ nodes with missing Caspr^+^ paranodes bilaterally (as a ratio of all Nfasc^+^ nodes; Fig. 1B; p = 0.0007), and an increase in Nfasc^+^ nodes with only a unilateral Caspr^+^ paranode (as a ratio of all Nfasc^+^ nodes; Fig. 1C; p = 0.0359), when compared to WT. Knock-down of TDP-43^Q331K^ in Schwann cells rescued these phenotypes, with the ratio of missing bilateral Caspr^+^ paranodes to all Nfasc^+^ (Fig. 1B; p = 0.0005) being comparable to that seen in WT mice. Analysis of nodal morphology (Fig. 1D) revealed a reduction in mean node length in TDP-43 mice compared to WT (Fig. 1E; p = 0.0101). This deficit was rescued in the P0-cre/TDP-43 mice (p = 0.0221), a pattern that was also reflected in the distribution of individual node lengths across genotypes (Fig. 1F). There were no differences between genotypes with respect to mean Nfasc^+^ node area, despite a trend towards a decrease in TDP-43 compared to WT (Fig. 1G; p = 0.0601), or mean area of Caspr^+^ paranodes (Fig. 1H). These pathologies were present in both motor and sensory nerves, (Supplementary Fig. 2) and the intensities of Caspr and Nfasc proteins remained unchanged between genotypes or between nerve type (Supplementary Fig. 3).

**Figure 1.**
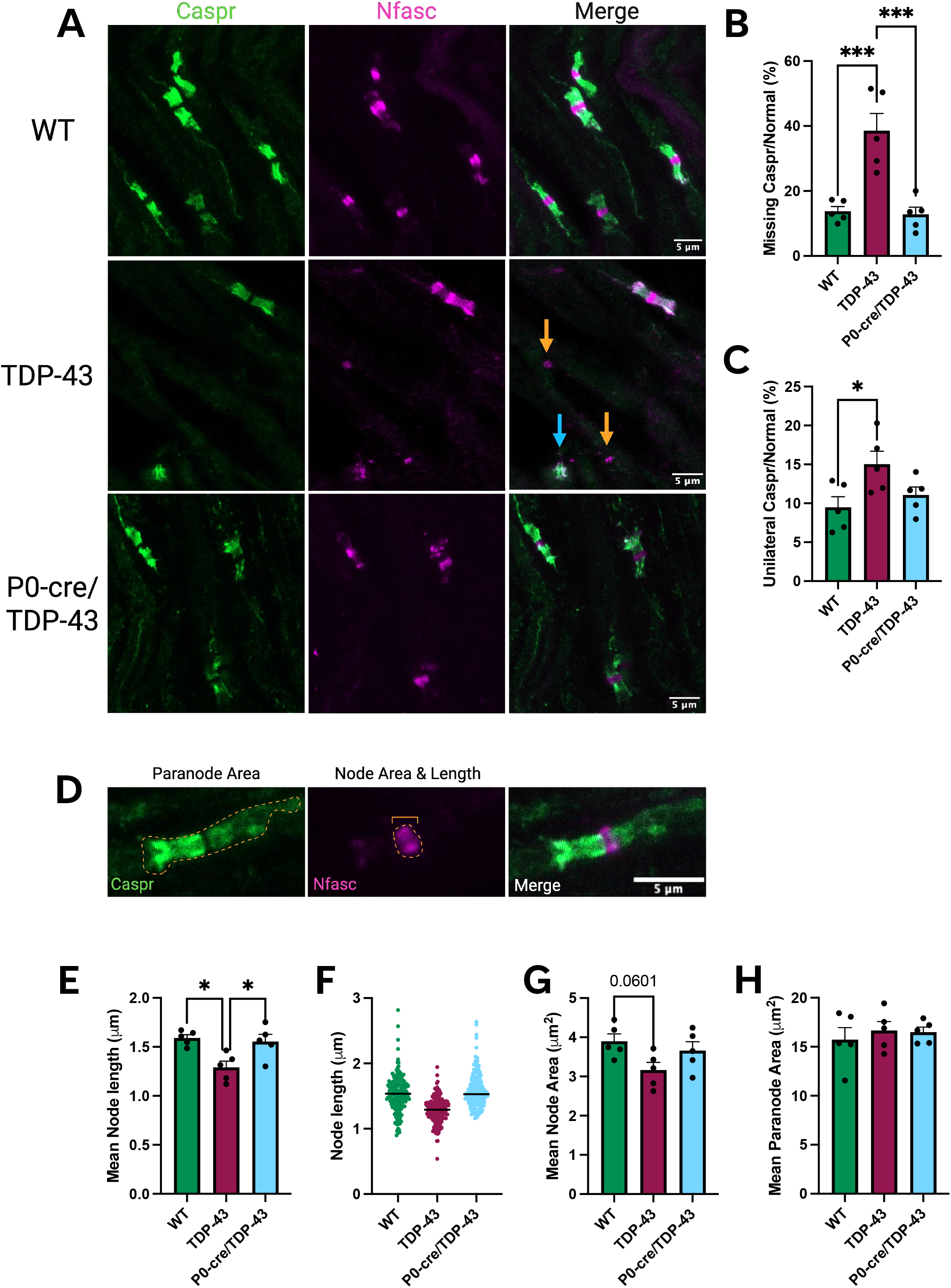
Node and paranode structures in the sciatic nerves of 10 mo mice. **A.** Representative images of nodes and paranodes within WT, TDP-43, and P0-cre/TDP-43 sciatic nerves immunolabelled for contactin-associated protein (Caspr; green) and neurofascin (Nfasc; magenta). Nodes of Ranvier missing paranodes bilaterally (orange arrows) or unilaterally (blue arrow) were observed; scale bars 5 µm. **B.** The density of missing paranodes as a ratio to normal Node of Ranvier assembly was increased in TDP-43 mice compared to WT (p = 0.0007) and was rescued in the P0-cre/TDP-43 mice compared to TDP-43 (p = 0.0005). **C.** The ratio of unilateral Caspr^+^ paranodes to normal was also significantly increased in the TDP-43 mice compared to WT (p = 0.0359)**. D.** Representative images of a paranode with the parameters for measuring paranode area with Caspr (green), and node area and length with Nfasc (magenta), scale bars 5 µm. **E**. Mean node length was significantly decreased in TDP-43 mice compared to WT (p = 0.0101), which was rescued in P0-cre/TDP-43 mice compared to TDP-43 (p = 0.0221). **F.** Frequency distribution of individual node lengths for each genotype. **G**. The mean node area exhibited a decreased trend for TDP-43 mice compared to WT (p = 0.0601), with P0-cre/TDP-43 mice unchanged from WT. **H.** There was no change between genotypes in mean paranode area. All data are presented as mean ± S.E.M, One-way ANOVA with Tukey’s *post hoc* test; *p < 0.05, **p < 0.01, ***p < 0.001, ****p < 0.0001; n = 5 mice per genotype, 100-150 nodes and paranodes analysed per animal.

### PNS Myelin Abnormalities in TDP-43 Mice Are Not Seen in P0-cre/TDP-43 Mice

Given the paranodal and nodal structural disruptions, we next characterised internode morphology, the myelinated segment of the axon. Utilizing Spectral Confocal Reflectance (SCoRe) microscopy, which uses the reflected laser light off compact, lipid-dense myelin, we quantified myelin reflectance and myelin density (Schain et al., 2014) (Fig. 2A). In sciatic nerves from 10 mo mice sectioned longitudinally, myelin reflectance was significantly higher in TDP-43 mice compared to WT (Fig. 2B; p = 0.0238). To understand changes occurring in the myelin on an ultrastructural level, we performed transmission electron microscopy (TEM) on transverse sections of sciatic nerves from 10 mo mice to measure g-ratio, which is the ratio of myelin thickness to axon diameter (Fig. 2C). The g-ratios (Fig. 2D; Supplementary Fig. 4A-D) and axon diameter frequencies (Fig. 2E) remained unchanged between genotypes. In previous work, we identified that increased myelin reflectance obtained from SCoRe, could be due to increased myelin degeneration, misfolded myelin, and immature myelin with no quantifiable change to g-ratio or axon density (Lewis et al., 2025), thus we wanted to identify potential ultrastructural deficits in the myelin. Assessment of myelin ultrastructure revealed the density of myelinated axons without apparent morphological deficit was significantly decreased in TDP-43 mice compared to WT (p = 0.0302) and P0-cre/TDP-43 (p = 0.0017; Supplementary Fig. 4E) mice. This was irrespective of there being no change to the total density of myelinated axons (Supplementary Fig. 4F). In TDP-43 mice, key myelin abnormalities were identified, including increased axons with misfolded myelin (p = 0.0012; Fig. 2J), non-compact myelin (p = 0.0320; Fig. 2K), and axons with axonal or myelin degeneration (p < 0.0001; Fig. 2L), compared to WT. All three abnormalities were rescued in P0-cre/TDP-43 mice (p = 0.0353, p = 0.0224 and p < 0.0001, respectively). Overall, we identified key myelin phenotypes in TDP-43 mice which were all ameliorated in P0-cre/TDP-43 mice.

**Figure 2.**
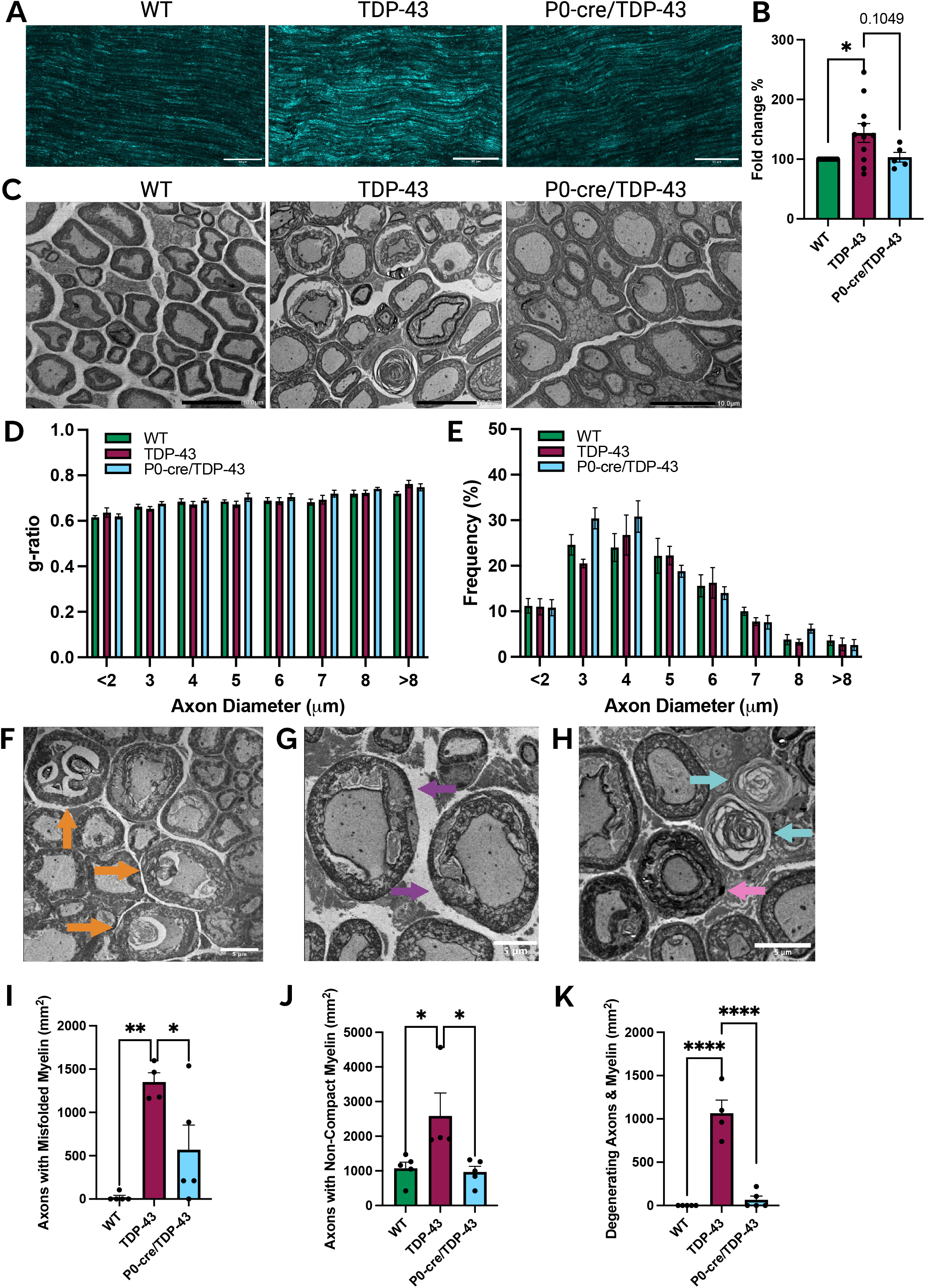
Myelin analyses of sciatic nerves of 10 mo mice. **A.** Representative Spectral Confocal Reflectance (SCoRe) Microscopy images of WT, TDP-43, and P0-cre/TDP-43 sciatic nerves, scale bars 50 µm. **B.** TDP-43 mice displayed an increased SCoRe signal, represented as a fold change comparative to WT (p = 0.0238), with P0-cre/TDP43 showing a trend of a rescue compared to TDP-43 (p = 0.1049). **C.** Representative images of electron microscopy (EM) of 10 mo sciatic nerve cross sections, scale bars 10 µm. G-ratios were unchanged between genotypes, independent of axon diameter (**D**). There were no differences in the distributions of axon diameters sampled in the sciatic nerves (**E**). Representative images of pathology within TDP-43 sciatic nerves displaying myelin with abnormal infoldings and myelination within the axonal space (**F**; orange arrows), axons with non-compact myelin (**G;** purple arrows), axonal degeneration (**H**; blue arrows), and myelin degeneration (**H**; pink arrow), scale bars 5 µm. **I.** TDP-43 mice showed increased densities of axons with misfolded myelin compared to WT (p = 0.0012) and P0-cre/TDP-43 (p = 0.0353), axons with non-compact myelin compared to WT (**J**; p = 0.0320) and P0-cre/TDP-43 (p = 0.0224), and axons with evidence of degeneration compared to WT (**K**; p < 0.0001) and P0-cre/TDP-43 (p < 0.0001). All data are presented as mean ± S.E.M or frequency distributions, One-way ANOVA with Tukey’s *post hoc* test; *p < 0.05, **p < 0.01, ***p < 0.001, ****p < 0.0001; n = 4-11 mice per genotype. G-ratio calculated from 100-150 axons per animal.

### Both TDP-43 and P0-cre/TDP-43 Mice Experience Motor Behaviour Deficits

Since healthy Schwann cells were sufficient to rescue nodal and paranodal histology and myelin pathology in P0-cre/TDP-43 compared to TDP-43 mice, we aimed to assess if these structural changes also rescued functional outputs in mice. WT, TDP-43, and P0-cre/TDP-43 were weighed monthly throughout the disease course (1 mo – 10 mo), where it was established that P0-cre/TDP-43 mice consistently weighed more than WT and sometimes TDP-43 (Supplementary Fig. 5A, B). Since weight can impact motor behavioural performance [41], all behavioural data were corrected for individual mouse weight at each time point (non-corrected data in Supplementary Fig. 5C-F). Hind limb grip strength and rotarod testing were performed fortnightly from 2 mo to 10 mo. Longitudinal analyses of the rotarod test showed that the latency to fall in TDP-43 and P0-cre/TDP-43 mice was shorter than WT mice (Fig. 3A; p < 0.0001). At 10 mo, both TDP-43 and P0-cre/TDP-43 mice fell significantly faster than WT mice (Fig. 3B; p < 0.0001 and p < 0.0001, respectively). There were significant decreases in hind limb grip strength in both TDP-43 and P0-cre/TDP-43 mice (Fig. 3C; p = 0.0054) when compared to WT mice. At 10 mo, P0-cre/TDP-43 mice performed significantly worse than WT (Fig. 3D; p = 0.0005) and TDP-43 showed a decreased trend compared to WT (p = 0.0998). Thus, both TDP-43 and P0-cre/TDP-43 mice exhibited worsened motor behaviour throughout the disease course compared to WT.

**Figure 3.**
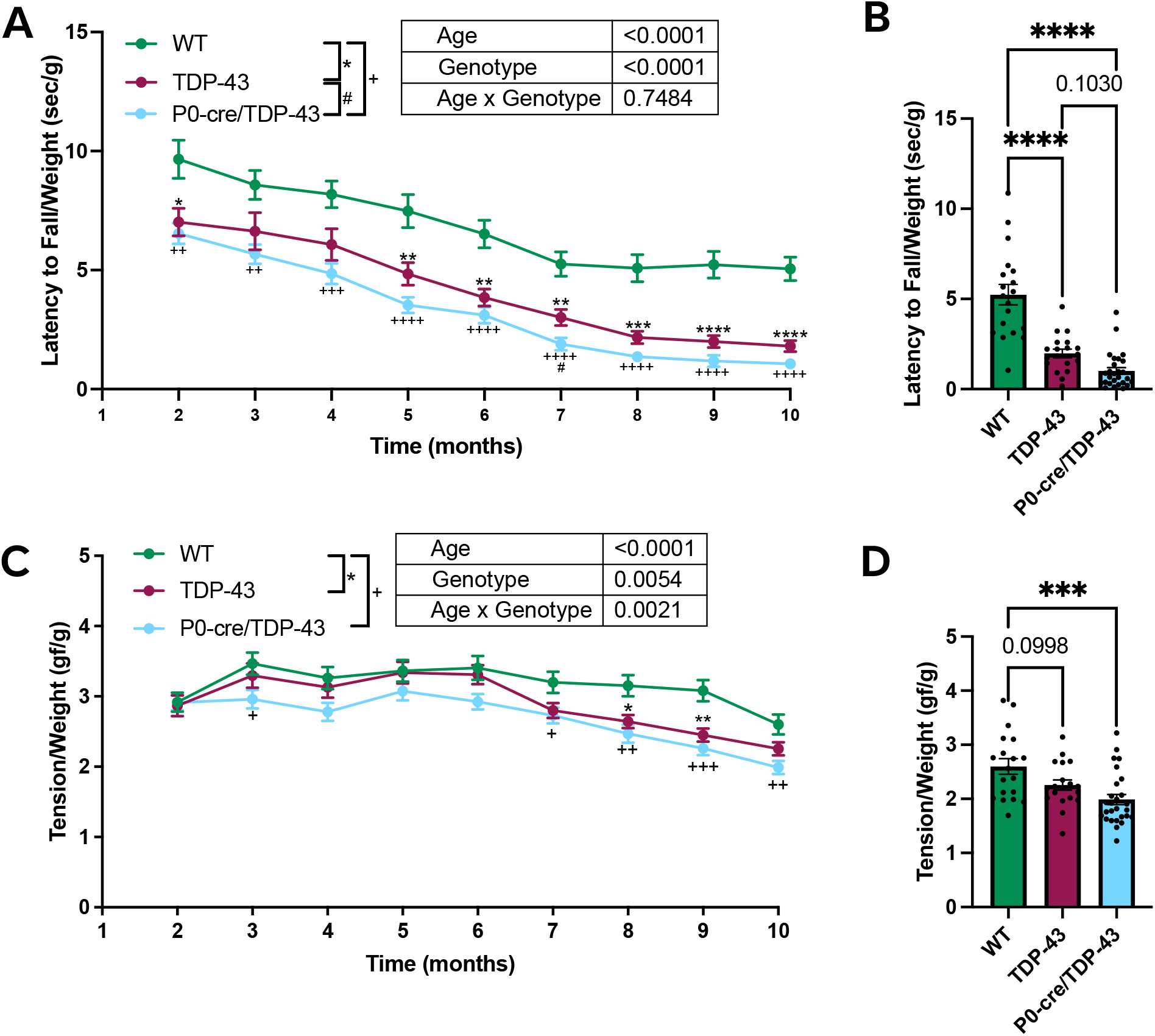
Weight normalised hind-limb grip strength and the rotarod walking test for WT, TDP-43, and P0-cre/TDP-43 mice. **A.** Weight normalised rotarod latency to fall was significantly reduced for TDP-43 and P0-cre/TDP-43 mice compared to WT (p < 0.0001), with TDP-43 falling faster than WT at 2, 5, 6, 7, 8, 9, and 10. P0-cre/TDP-43 performed worse than WT at every time point; 2 mo – 10 mo and fell faster than TDP-43 at 7 mo. **B.** At 10 mo, TDP-43 and P0-cre/TDP-43 fell faster on the rotarod test than WT (p < 0.0001 and p < 0.0001, respectively) and there was a trend for faster fall time for P0-cre/TDP-43 compared to TDP-43 (p = 0.1030). **C.** Weight normalised hind limb grip strength showed both TDP-43 and P0-cre/TDP-43 having reduced grip strength (p = 0.0054), with TDP-43 performing worse than WT at 7 and 8 mo and P0-cre/TDP-43 scoring lower than WT at 3, 7, 8, 9, and 10 mo. **D.** At 10 mo (10 mo), P0-cre/TDP-43 mice had significantly reduced grip strength compared to WT (p = 0.0005) and TDP-43 showed a decreased trend compared to WT (p = 0.0998). All data are presented as mean ± S.E.M, Mixed effects analysis with Tukey’s post-hoc analyses for line graphs, One-Way ANOVAs for 10 mo analyses, n = 19-28 (12 F; 8-16 M) per genotype, *p<0.05, **p<0.01, ***p<0.001, ****p<0.0001.

### Sciatic Nerve Conduction Deficits Are Present in both TDP-43 and P0-cre/TDP-43 mice

Alterations to nodal and paranodal proteins and node size can directly influence neuronal conduction velocity (Arancibia-Carcamo et al., 2017; Babbs & Shi, 2013; Bhat et al., 2001; Chang et al., 2021; Hasegawa et al., 1988; Rosenbluth & Bobrowski-Khoury, 2014). We therefore conducted sciatic nerve electrophysiology to assess if nerve conduction was impacted by Schwann cell specific removal of mutant TDP-43. While TDP-43 mice have a well characterised compound muscle action potential (CMAP) deficit (Arnold et al., 2013; Watkins et al., 2021), conduction velocity has not been investigated. However, there are also conflicting reports on the level of motor neuron loss in this mouse model, with some research suggesting no lower motor neuron death even at 24 mo (Watkins et al., 2021). Hence, we first ascertained the degree of neurodegeneration prior to assessing the impact on function.

To quantify lower motor neuron loss, we delivered Fluorogold via intraperitoneal injection, which is a retrograde tracer taken up at the neuromuscular junction and deposited into motor neuronal cell bodies (Anderson & Edwards, 1994; Mohan et al., 2014; R Mohan et al., 2015). This ensured the evaluation of changes to motor axon density independent of changes to cell soma density. We assessed motor neuron density by co-labelling ChAT^+^ and Fluorogold^+^ motor neurons in the ventral horn grey matter of the lumbar spinal cord of 10 mo WT and TDP-43 mice (Supplementary Fig. 6A). We observed no differences in the densities of intact motor neurons between WT and TDP-43 mice (Supplementary Fig. 6B-D), thus confirming no overt loss of lower motor neurons in TDP-43 mice. Due to there being no difference, we did not perform this assessment in P0-cre/TDP-43 mice.

Having established that any alterations in sciatic nerve conduction would not be skewed by motor neuron loss, we proceeded to perform sciatic nerve electrophysiology on 10 mo WT, TDP-43, and P0-cre/TDP-43 mice. We utilised an array of three stimulating electrodes, spaced evenly 2 mm apart, to allow measurements of sciatic nerve motor activity. A recording electrode positioned on the gastrocnemius muscle recorded compound muscle action potential (CMAP) waves, or M-waves, which were averaged to provide a single M-wave per animal, represented as the average waveform for each genotype (Fig. 4A). Mean CMAP amplitude was significantly decreased in both TDP-43 and P0-cre/TDP-43 mice compared to WT (Fig. 4B; p = 0.0072 and p = 0.0022, respectively). The mean onset latency, the time taken between nerve stimulation and the beginning of the averaged M-wave, was significantly delayed for both TDP-43 and P0-cre/TDP-43 compared to WT (Fig. 4C; p = 0.0265 and p = 0.0170, respectively), while the duration of the M-wave remained unchanged between the three genotypes (Fig. 4D). The mean area under the curve, accounting for peak and duration of CMAP, was significantly decreased for TDP-43 and P0-cre/TDP-43 compared to WT (Fig. 4E; p = 0.0109 and 0.0041, respectively).

**Figure 4.**
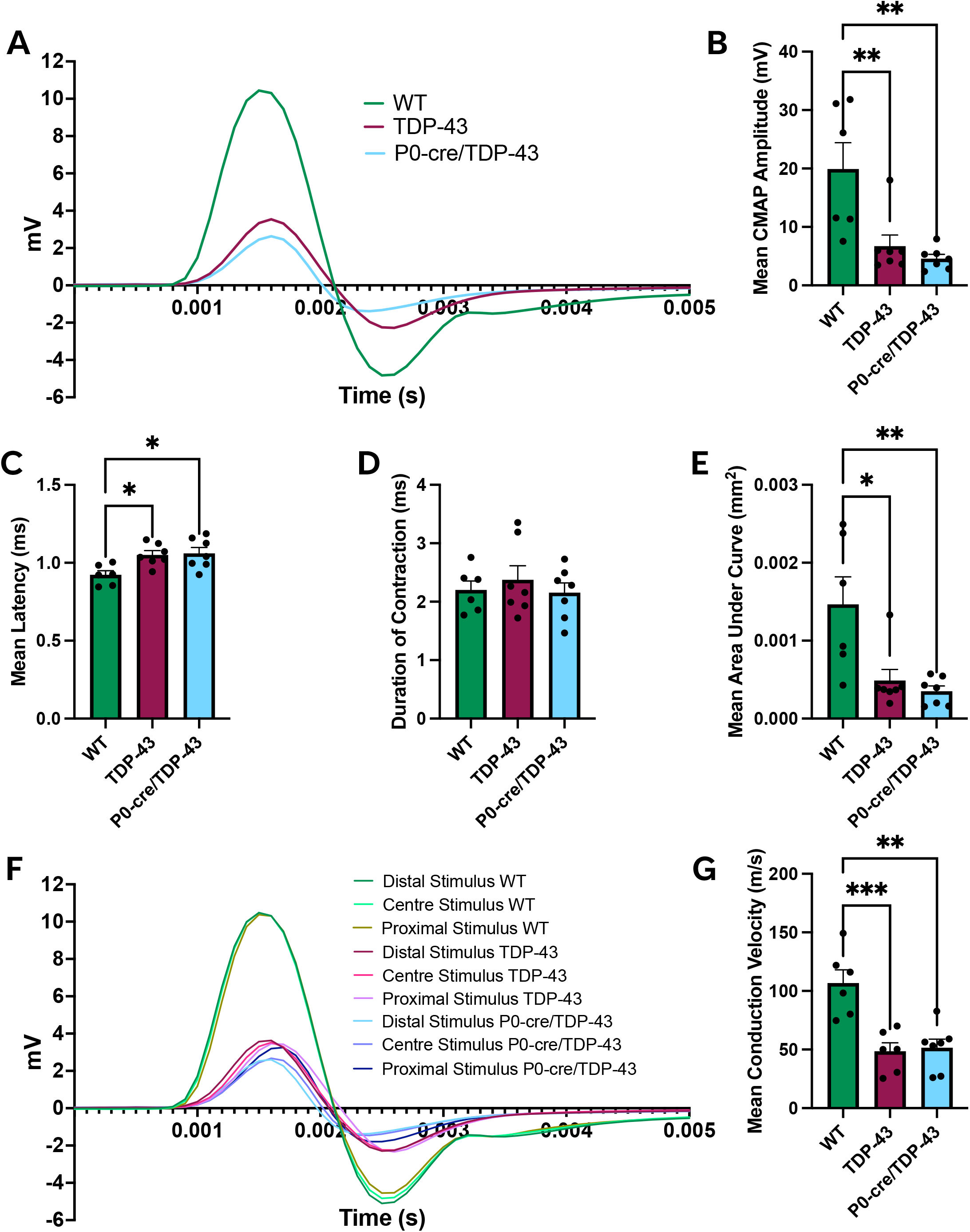
*In vivo* sciatic nerve electrophysiology recordings on 10 mo WT, TDP-43, P0-cre/TDP-43 mice. **A.** Average M-waves of all three stimuli for each genotype. **B.** Mean Compound Muscle Action Potential (CMAP) amplitude showed a significant decrease for both TDP-43 and P0-cre/TDP-43 compared to WT (p = 0.0072 and p = 0.0022, respectively). **C.** Mean onset latency, i.e. the time between stimulation and the beginning of the CMAP, was significantly delayed for both TDP-43 and P0-cre/TDP-43 compared to WT (p = 0.0265 and p = 0.0170, respectively). **D.** The mean duration of the M-wave was unchanged between genotypes. **E.** The mean area under the curve was significantly decreased for both TDP-43 and P0-cre/TDP-43 compared to WT (p = 0.0109 and 0.0041, respectively). **F.** Mean M-waves for each electrode stimulus for each genotype, showing the spread in each waveform. **G.** Mean sciatic nerve conduction velocity was significantly decreased in both TDP-43 and P0-cre/TDP-43 compared to WT (p = 0.0009 and p = 0.0010, respectively). All data are presented as mean ± S.E.M, Ordinary One-Way ANOVA, with Tukey’s post-hoc analyses; n = 6-7 (2-3 F; 4 M) per genotype; *p<0.05, **p<0.01, ***p<0.001.

While CMAP amplitude, onset latency, and area under the curve provide meaningful information regarding the stimulation and muscle action potential response, the delays and reductions observed in the TDP-43 and P0-cre/TDP-43 mice could be due to neuromuscular deficits, or pathology of terminal Schwann cells, a subtype of Schwann cell that forms caps at the axon terminal and neuromuscular junction and allows for the correct formation of the neuromuscular junction (Darabid et al., 2013; Kang et al., 2014; Smith et al., 2013). To rule neuromuscular pathologies out as the only contributor to these electrophysiology deficits, we measured M-waves from each of the three stimulating electrodes. Knowing the distance and calculating the time between each stimulus, we could accurately examine sciatic nerve conduction time between the first and last stimulus, with no neuromuscular junction or muscular influence (Fig. 4F). Indeed, there was a significant reduction in sciatic nerve conduction velocity in TDP-43 and P0-cre/TDP-43 compared to WT (Fig. 4G; p = 0.0009 and p = 0.0010, respectively). Taken together, both the TDP-43 and P0-cre/TDP-43 mice exhibit worsened electrophysiological deficit, independent of neuromuscular junction involvement.

After revealing no functional rescue in P0-cre/TDP-43 mice compared to TDP-43 at a behavioural and circuitry level, we postulated that functional rescue may be at a sub-cellular level. Retrograde axonal transport deficits have previously been implicated in motor neuron dysfunction across several ALS mouse models (Sleigh, Tosolini, Gordon, et al., 2020; Tosolini et al., 2024; Tosolini et al., 2022), however axonal transport has not been assessed in the TDP-43^Q331K^ mouse. Of interest, signalling endosome and mitochondrial transport can slow at the nodes of Ranvier forming bottlenecks (Tosolini et al., 2024), raising the possibility that nodal architecture may influence signalling endosome dynamics, and therefore be affected by the rescue of nodal pathology in our P0-cre/TDP-43 mice. To investigate this, we performed *in vivo* axonal transport imaging on anaesthetized 10 mo WT, TDP-43, and P0-cre/TDP-43 mice (Supplementary Fig. 7A; Supplementary Video 1). However, no differences were detected in any retrograde axonal transport dynamics (Supplementary Fig. 7B-G). These findings suggest that retrograde signalling endosome transport remains preserved in TDP-43^Q331K^ mice and is therefore unlikely to contribute substantially to the neuronal or myelin abnormalities observed in this model.

### Büngner Schwann Cells are Activated in Both TDP-43 and P0-cre/TDP-43 mice

The lack of functional improvement in TDP-43 mice following rescue in nodal and paranodal assembly and myelin architecture in P0-cre/TDP-43 mice led us to question the involvement of other subtypes of Schwann cells. Under the influence of c-Jun, myelinating and non-myelinating Schwann cells undergo transformation into Büngner Schwann cells (Arthur-Farraj et al., 2012). This unique type of Schwann cell has previously been implicated in ALS rodent models (Cabeza-Fernández et al., 2024; Trias et al., 2020), whereby the selective overexpression of c-Jun in Schwann cells of SOD1^G93A^ led to worsened motor performance, compared to regular SOD1^G93A^ mice (Cabeza-Fernández et al., 2024). While Büngner Schwann cells can facilitate axon repair after acute injury or degeneration (Jessen & Mirsky, 2016), chronic activation of Büngner Schwann cells can cause detrimental inflammatory effects (Berner et al., 2022; Gomez-Sanchez et al., 2015; Hartlehnert et al., 2017; Wagner & Myers, 1996), potentially contributing to the poorer functional outputs observed in the P0-cre/TDP-43 mice. Since Büngner Schwann cells arise from reprogrammed myelinating and non-myelination Schwann cells, they also express P0. Thus, we sought to establish whether seemingly healthy Büngner Schwann cells in the P0-cre/TDP-43 mice were still sensitive to the surrounding pathological environment. Second to this, we sought to determine whether this sensitivity may underpin the divergent phenomena of nodal and myelin normalization, and persistent motor and electrophysiological deficit. We immunolabelled 10 mo sciatic nerve sections for c-Jun, and glial fibrillary acidic protein (GFAP), which is produced by Büngner Schwann cells during and after reprogramming (Kobayashi et al., 1986), alongside the pan Schwann cell transcription factor SOX10 (Fig. 5A). The ratios of c-Jun^+^ Schwann cells or GFAP^+^ Schwann cells to all SOX10^+^ Schwann cells were significantly increased in TDP-43 and P0-cre/TDP-43 mice compared to WT mice (Fig. 5B, p = 0.0013 and p = 0.0383, respectively; Fig. 5C, p < 0.0001 and p < 0.0001, respectively). The ratio of c-Jun^+^ Schwann cells with GFAP, as a proportion of all SOX10^+^ Schwann cells was also increased for TDP-43 and P0-cre/TDP-43 compared to WT (Fig. 5D, p < 0.0001 and p = 0.0005, respectively). Since Büngner Schwann cells recruit macrophages (Berner et al., 2022; Cabeza-Fernández et al., 2024; Gomez-Sanchez et al., 2015; Hartlehnert et al., 2017), which can exacerbate nerve inflammation and worsen disease progression in ALS models (Cabeza-Fernández et al., 2024; Trias et al., 2018; Trias et al., 2020), we hypothesized that the Büngner Schwann cell transdifferentiation correlated with heightened inflammation. Indeed, we identified significant immune cell infiltration. To ascertain altered macrophagic density, we immunolabelled sciatic nerves for both Iba1, a pan macrophagic marker, and CD11b which marks activated macrophages (Robinson et al., 1986) (Fig. 5E) and its expression is linked to worsened ALS disease progression (Graber et al., 2010; Van Dyke et al., 2016). The density of Iba^+^ macrophages was significantly increased in both TDP-43 and P0-cre/TDP-43 mice (Fig. 5F; p = 0.0120 and p = 0.0100, respectively) and Iba1^+^CD11b^+^ activated macrophages was also increased in TDP-43 and P0-cre/TDP-43 mice (Fig. 5G; p < 0.0001 and p < 0.0001, respectively), reflecting significant macrophage infiltration and activation. Therefore, despite the rescue in myelinating Schwann cell pathology, the recruitment of Büngner Schwann cells, and the inflammatory cascade associated with their activation (Berner et al., 2022; Gomez-Sanchez et al., 2015; Hartlehnert et al., 2017; Wagner & Myers, 1996), may contribute to the overall functional deficits observed in P0-cre/TDP-43 mice.

**Figure 5.**
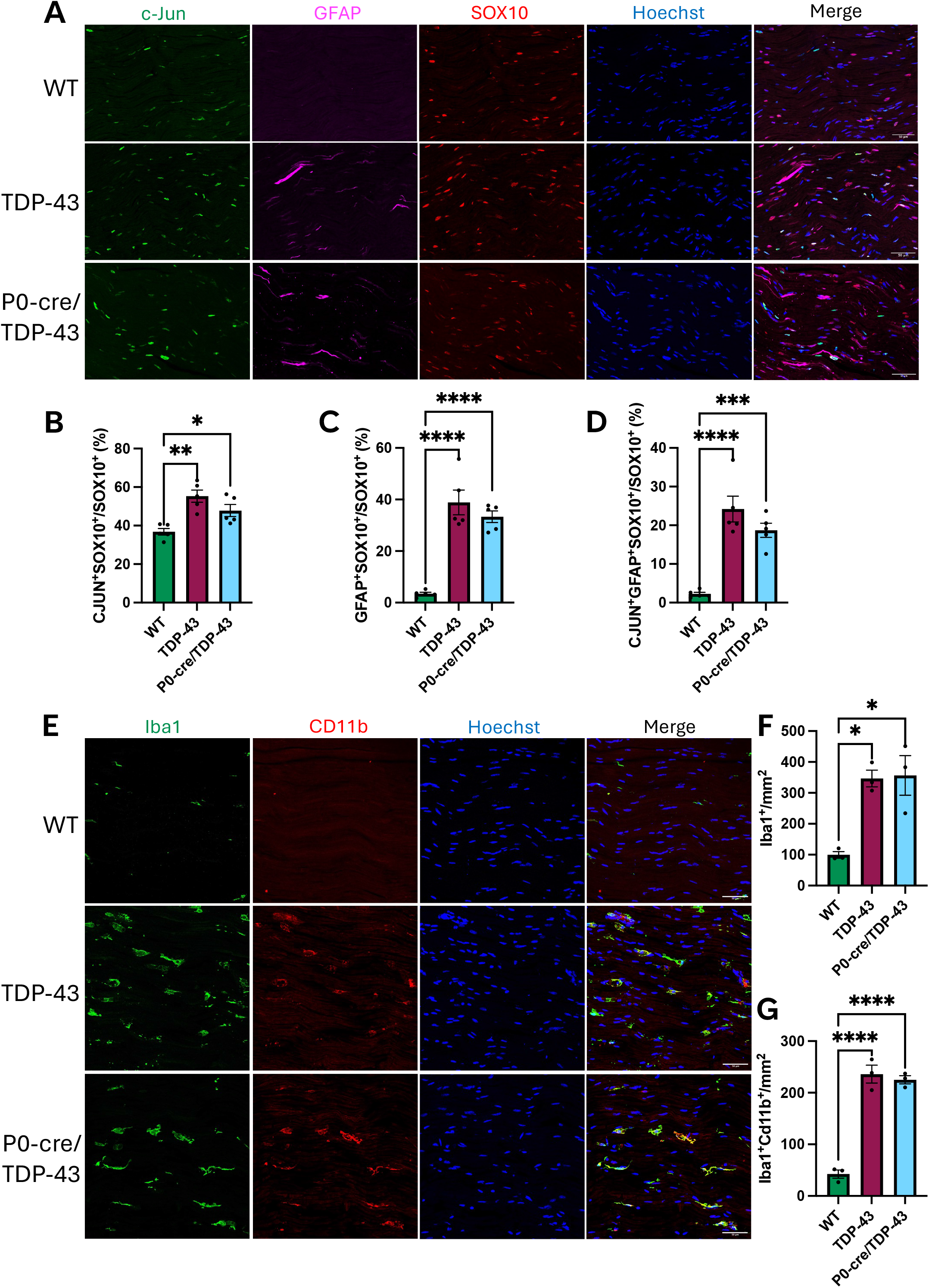
Repair Schwann cell and macrophagic activation in sciatic nerves of 10 mo mice. **A.** Representative images of WT, TDP-43, and P0-cre/TDP-43 sciatic nerves marking activated repair Schwann cells (c-Jun, green; GFAP, magenta), all Schwann cells (SOX10, red), and all nucleated cells (Hoechst, blue), scale bars 50 µm. B. The ratio of co-labelled c-Jun^+^SOX10^+^ Schwann cells to all SOX10^+^ Schwann cells was significantly increased in both TDP-43 and P0-cre/TDP-43 mice compared to WT (p = 0.0013 and p = 0.0383, respectively) and was unchanged between TDP-43 and P0-cre/TDP-43. **C.** The ratio of GFAP^+^SOX10^+^ Schwann cells to all SOX10^+^ Schwann cells was also significantly increased in both TDP-43 and P0-cre/TDP-43 mice compared to WT (p < 0.0001 and p < 0.0001). **D.** The ratio of c-Jun^+^GFAP^+^SOX10^+^ Schwann cells to all SOX10^+^ Schwann cells was increased for TDP-43 and P0-cre/TDP-43 compared to WT (p < 0.0001 and p = 0.0005, respectively). **E.** Representative images of WT, TDP-43, and P0-cre/TDP-43 sciatic nerves marking Iba1^+^ (green) and CD11b^+^ (red) macrophages, plus all nucleated cells (Hoechst, blue). **F.** TDP-43 and P0-cre/TDP-43 mice both expressed significantly higher densities of Iba1^+^ cells compared to WT (p = 0.0120 and p = 0.0100, respectively), as well as more Iba1^+^Cd11b^+^ cells per mm^2^ (**G**; p < 0.0001 and p < 0.0001, respectively). Mean ± S.E.M, *p < 0.05, **p < 0.01; ***p < 0.001; ****p < 0.0001; One-way ANOVA with Tukey’s post hoc test; n = 3-5 mice per genotype.

## Discussion

Schwann cells are essential for maintaining PNS homeostasis through modulating nerve conduction, providing energy to the neuron (Belin et al., 2017; Bouçanova et al., 2021; Deck et al., 2022; Nave & Werner, 2021; Uncini et al., 2024), immunological functions (Berner et al., 2022; Dubový et al., 2014; Martini et al., 2008), influencing synaptic transmission (Hyung et al., 2018; Reddy et al., 2003; Robitaille, 1995; Todd et al., 2010), guiding the formation of the neuromuscular junction (Darabid et al., 2013; Kang et al., 2014; Smith et al., 2013), and driving protein complex formation at and around the nodes of Ranvier (Chang et al., 2021; Eshed-Eisenbach et al., 2023; Poliak & Peles, 2003). Given their importance in supporting neuronal health, if Schwann cells are dysfunctional in ALS, their pathology may contribute to, and exacerbate, neuronal dysfunction. This is the first study to investigate Schwann cells and PNS myelin in the TDP-43^Q331K^ mouse model. In TDP-43^Q331K^ mice, we found changes to node and paranode structure, and disrupted myelination, and when the TDP-43^Q331K^ mutation was selectively removed from Schwann cells, these phenotypes were rescued. Despite the histological rescue, there was no improvement in motor behaviour or electrophysiological properties in P0-cre/TDP-43 mice. This, at least in part, could be attributed to the activation of Büngner Schwann cells, which are known to exacerbate inflammation (Berner et al., 2022; Gomez-Sanchez et al., 2015; Hartlehnert et al., 2017; Wagner & Myers, 1996). While rescuing myelinating Schwann cell TDP-43 proteinopathy indeed showed protective benefits to myelin pathology in this mouse, the sole rescue of myelin may not be sufficient to rescue lower motor neuron deficits in TDP-43 associated ALS.

This study assessed peripheral myelination in the TDP-43^Q331K^ transgenic mouse model. Previous studies have shown that selective deletion of endogenous *TARDBP* in Schwann cells in mice caused profound structural changes to PNS paranodes, slowed nerve conduction, and caused motor behavioural deficits (Chang et al., 2021). What has remained unclear is whether such phenotypes are replicated in a mouse over-expressing a mutant human *TARDBP* transgene. We show that the TDP-43^Q331K^ mouse exhibits loss of paranodal proteins, slowed conduction velocity, and poorer motor performance. While we found no alterations in g-ratio, we observed an increase in myelin reflectance and evidence of misfolded, non-compact, and degenerating myelin, which were similarly found in spinal cord myelin in this TDP-43^Q331K^ mouse model (Lewis et al., 2025). Although the lack of change in g-ratio aligns with what has been shown previously (Chang et al., 2021), it contrasts reports of no ultrastructural myelin pathology in mice with Schwann cell-specific *TARDBP* knock down (Chang et al., 2021). This raises the question as to whether structural abnormalities in Schwann cells results from a gain of function pathology, rather than loss of function, or whether they arise in response to the surrounding diseased environment. A recent study assessing global *C9orf72* deletion in SOD1 ALS mice revealed increased sciatic nerve g-ratio, hence thinner myelin, when compared to WT (Sironi et al., 2025), suggesting that myelin changes may contribute to ALS, irrespective of genetics. In P0-cre/TDP-43 mice, rescue of nodal, paranodal, and myelin abnormalities was not sufficient to attenuate functional phenotypes. These data indicate that motor and electrophysiological deficits in TDP-43 mice are unlikely to be driven by myelinating Schwann cells. To determine the intrinsic dysfunction in Schwann cells in ALS, a future study to selectively overexpress the human mutant TDP-43 protein in Schwann cells alone and assess any dysfunction may be of value. Understanding how the loss and gain of function of TDP-43 impact Schwann cells and PNS myelin is of key interest in ALS and will help establish how these cells are contributing to the disease.

Despite the rescue in most major histological phenotypes, P0-cre/TDP-43 mice did not exhibit any functional rescue. We showed worsened motor performance in TDP-43 mice, compared to WT, as has been characterised previously (Arnold et al., 2013; Lewis et al., 2025). We predicted the motor deficits would be rescued in our P0-cre/TDP-43 mice; however, this was not observed. Regarding sciatic nerve electrophysiology, as expected, the TDP-43 mice had worsened CMAP amplitude, onset latency, and conduction velocity, compared to WT mice. The onset latency and conduction velocity have not been previously assessed in TDP-43^Q331K^ mice, while our CMAP data align with those published previously (Arnold et al., 2013; Watkins et al., 2021). We also confirmed there was no evidence of motor neuron loss in our cohort of TDP-43 mice, which had also been found previously in this model (Mitchell et al., 2015). Regardless, the electrophysiological deficits observed cannot be attributed to fewer available motor neurons firing. Since we ascertained the conduction velocity deficits by recording the velocity along different points of the axon, it rules out deficits in neuromuscular junction function (and potential influence of terminal Schwann cells) as being the only contributor to the impairment. Surprisingly, selectively deleting the mutant *TDP-43* transgene from Schwann cells alone did not rescue any of these electrophysiological phenotypes. Therefore, the rescue in nodal and paranodal structure in the P0-cre/TDP-43 mice was not sufficient to ameliorate electrophysiological deficits, meaning dysfunction in nerve conduction in the TDP-43 and P0-cre/TDP-43 mice is not myelin driven. Given the direct correlation between nodal assembly and nerve conduction (Arancibia-Carcamo et al., 2017; Chang et al., 2021), an alternate explanation was required to ascertain why sciatic nerve functions were not similarly rescued. One such avenue was to investigate the Büngner Schwann cells, a specialized type of Schwann cell that reprograms from myelinating and non-myelinating Schwann cells in response to inflammation and neuronal damage (Jessen & Mirsky, 2016; Kyriakis et al., 1994). We established that there was significant recruitment of Büngner Schwann cells in both TDP-43 and P0-cre/TDP-43 mice. Previously, in SOD1^G93A^ mice, the selective overexpression of c-Jun in Schwann cells caused worsened motor performance and macrophagic immune infiltration (Cabeza-Fernández et al., 2024), aligning with what we observed in both TDP-43 and P0-cre/TDP-43 mice. It is well established that Büngner Schwann cells release pro- and anti-inflammatory cytokines and recruit macrophages for the purpose of clearing debris and assisting in the axonal repair process (Berner et al., 2022; Gomez-Sanchez et al., 2015; Hartlehnert et al., 2017; Wagner & Myers, 1996), however, chronic inflammation and chronic activation of Büngner Schwann cells can result in hyperinflammation and reduced capacity for these cells to initiate neuronal repair (Büttner et al., 2018). Moreover, inflammatory signals, such as TNF-α expression, have been shown to influence nerve conduction, whereby increased pro-inflammatory cytokine expression decreased nerve conduction capacity (Allison et al., 2014; Davies et al., 2006; Magrinelli et al., 2015). Since many ALS patients experience PNS inflammation, often chronically (Lu et al., 2016; Schreiber et al., 2019), and ALS animal models also exhibit PNS inflammation (Chiu et al., 2009; Sironi et al., 2025; Trias et al., 2018), understanding the contributions of Büngner Schwann cells to this immune response in ALS is essential. We postulate that activation of Büngner Schwann cells, and the associated immune cell recruitment, is contributing to dysfunction in TDP-43 and P0-cre/TDP-43 mice and that preserving myelinating Schwann cell identify may offer neuroprotective effects. Ultimately, our findings highlight the importance of not treating all Schwann cell subtypes the same and studying each subtype and their contribution to ALS pathogenesis.

A major strength of this study is the integrated evaluation of PNS function through peripheral nerve electrophysiology, CMAP analysis, and *in vivo* axonal transport measurements. Regardless, it is important to acknowledge some of the limitations of this study. The overexpression of human TDP-43 in the TDP-43^Q331K^ mice is accompanied by a modest reduction in the endogenous mouse TDP-43 protein and mRNA expression (Arnold et al., 2013) and expression levels may have skewed following selective deletion of the human transgene. Moreover, there was a significant weight gain phenotype observed in the female TDP-43 and P0-cre/TDP-43 mice that was not identified in the males, which is a known phenotype of this TDP-43 mouse model (Watkins et al., 2021). While gut phenotypes in general are present in other TDP-43 mouse models (Esmaeili et al., 2013; Hatzipetros et al., 2014), the use of alternative diets may ameliorate these limitations (Coughlan et al., 2016; Herdewyn et al., 2014). Thus, the motor behaviour data was normalised by individual mouse weight to account for this weight gain, and only male mice were examined for histology.

This is the first study to examine Schwann cell function in the TDP-43^Q331K^ mouse model of ALS. In 10 mo mice, we identified clear pathology at the nodes of Ranvier in TDP-43 mice and altered myelin pathology, which were ameliorated in P0-cre/TDP-43 mice. Surprisingly, these changes were insufficient to lead to functional improvements in P0-cre/TDP-43 mice, possibly due to the activation of inflammatory Büngner Schwann cells. These findings suggest that whilst therapeutically targeting Schwann cells may be sufficient to rescue phenotypes at a cellular level, it cannot overcome the contributing influence of inflammation and neuronal dysfunction. Future research should further stratify by Schwann cell subtype to investigate both their individual contributions to ALS, but also the synergistic properties of targeting both myelin and neuronal phenotypes.

## Supplemental Videos

**Supplementary Video 1. *In vivo* imaging of retrograde signaling endosomes along axons in a 10 mo WT mouse.** Signaling endosomes are labelled with tetanus toxin fragment (HcT-555; magenta) within GFP-AAV transduced axons (green). Frame interval: 1.09 s. Playback rate = 10 fps. Acquisition time = 5 min 26 s. Scale bar 10 μm.

## Materials and Methods

### Ethics Statement

All animal experiments adhered to the Australian National Health and Medical Research Council published Code of Practice and were approved by the Florey Institute of Neuroscience and Mental Health Animal Ethics Committee (Ethics numbers: 20-073-FINMH, 20-076-FINMH), and the University of Queensland’s Animal Ethics Committee (Ethics number: 2023/AE000649). Animals were housed in Tecniplast Sealsafe Plus GM500 cages (up to 6 mice per cage) with *ad libitum* access to food and water. Animals were monitored three times per week and weighed monthly.

### Generation of TDP-43^Q331K^/P0-cre Mice

TDP-43^Q331K^ mice (B6N.Cg-Tg(Prnp-TARDBP*Q331K)103Dwc/J, high expressing line 103, stock number 017933) and P0-cre mice (B6N.FVB-Tg(Mpz-cre)26Mes/J, stock number 017927) were purchased from The Jackson Laboratory (Bar Harbour, ME) and heterozygotes for both strains were crossed. This produced 4 separate genotypes: P0-cre^+^TDP-43Q331K^+^, P0-cre^+^TDP-43Q331K^-^, P0-cre^-^TDP-43Q331K^+^, and P0-cre^-^TDP-43Q331K^-^. Only P0-cre^+^TDP-43Q331K^+^ (P0-cre/TDP-43), P0-cre^-^TDP-43Q331K^+^ (TDP-43), and P0-cre^-^TDP-43Q331K^-^ (WT) mice were utilised for experiments as P0-cre^+^ mice have been characterised previously (Lobsiger et al., 2009). The TDP-43^Q331K^ human protein in the TDP-43 mouse is driven by the prion promotor, which has been identified in Schwann cells (Follet et al., 2002). Animals were genotyped either externally via TransnetYX or internally via the following protocol: lysis buffer (100mM Tris-HCl (pH 8.5), 5mM EDTA (pH 8.0), 200mM NaCl, 0.2% (w/v) SDS, and 100ug/mL proteinase K) added to tail clipping at 55 °C overnight in a heat bath. Proteinase K was deactivated in a heat bath for 99 °C for 10 min, then 1 min centrifuge at 13,000 rpm. Supernatant was then collected in separate tube, and equal volume isopropanol was added and mixed, then 13,000 rpm centrifuge for 10 min at room temperature, before discarding the supernatant and washing the pellet with 500uL of 70% Ethanol. The tube was centrifuged again at 13,000 rpm for 10 min at room temperature, the supernatant was then discarded and the tubes left to dry inverted over a tissue for 1 hr. The pellet was then re-dissolved in 50-100uL of Tris-EDTA buffer. For running the PCR genomic DNA was used at 50-100ng/ul, tested via a nanodrop, and 0.5-1ul was used per 20ul reaction (10uL quickload mastermix (New England Biolabs; M0271S), 1uL each primer, DNA (50-100ng), make up to 20ul with water). Primers were obtained from The Jackson Laboratory for B6N.Cg-Tg(Prnp-TARDBP*Q331K)103Dwc/J mice using protocol 27057 (Transgene Forward Primer 14748, Transgene Reverse Primer 14749, Internal Positive Control Forward Primer oIMR9020, Internal Positive Control Reverse Primer oIMR9021) and for B6N.FVB-Tg(Mpz-cre)26Mes/J using protocol 26258 (Transgene Forward Primer 14203, Transgene Reverse Primer oIMR9377, Internal Positive Control Forward Primer oIMR7338, Internal Positive Control Reverse Primer oIMR7339). PCR cycling was performed according to The Jackson Laboratory protocol 27057 (TDP-43) or 26258 (P0-cre). 1-2 % gels were run ∼80 V for 1 hr and imaged using the ChemiDoc™ MP (Bio-Rad). Each sample was tested for both TDP-43 and P0-cre protocols. WT, TDP-43 and P0-cre/TDP-43 mice were established as three colonies; one underwent behavioural testing with tissue collected at 10 mo and some undergoing nerve conduction tests at 10 mo (tissue not collected), the second were aged and tissue was collected at 2 mo, 5 mo, and 10, and the third underwent live axon transport imaging. Since the TDP-43 mouse exhibits a more predominant hind-limb pathology compared to fore-limb (Arnold et al., 2013), we focused on the sciatic nerve for all nerve analyses.

### Behavioural Assessment

Mice were weighed once per month from birth until sacrifice. Locomotor function was examined from 2 mo of age every 2 weeks via Rotarod (Ugo Basile) and Hind-limb grip strength (Able Scientific) tests. For Rotarod, animals were trained with two sessions of increasing speed from 2-40 RPM over 5 min, then rested for 10 min. Mice were assessed via 2-40 RPM increasing speed for a maximum of 5 min and the time at which they fell was recorded. This was performed twice per session, and an average was obtained per mouse. Hind-limb grip strength was measured through a T-Bar connected to a digital force gauge (Able Scientific), then holding the mice by their tail and lowering their hind-limbs onto the T-bar and pulling gently until their hind-limbs let go of the bar. 3 measurements per mouse per session were obtained, averaged, and were recorded in gram force. The same experimenter performed all hind-limb grip strength measurements to keep the pulling force constant.

### Immunoblotting

Mice were sacrificed via lethal injection (sodium pentobarbitone, 100 mg/kg, IP injection) then transcardially perfused with 0.1 M phosphate buffered saline (PBS). Fresh lumbar spinal cord and sciatic nerve tissue was collected at 2 mo and 10 mo from TDP-43 and P0-cre/TDP-43 mice. Sciatic nerves were dissected out between the sciatic notch and where the nerve bifurcates into the tibial and sural nerves and snap frozen. Tissues were thawed and homogenized in RIPA lysis buffer (50 mM Tris-Cl, pH 7.4, 150 mM NaCl, 1% TX-100, 1% sodium deoxycholate (Sigma), 0.1% SDS (AMRESCO), 1% protease inhibitor cocktail (Sigma P8340), 50 mM NaF, and 10 mM Na_3_VO_4_) and pulse sonicated at 50% output for 5 s bursts. Samples were then stored on ice for 20 min, then centrifuged at 4 °C at 13,000 RPM for 20 min. Supernatant was collected and stored in - 80 °C. Protein concentration in each sample was quantified using a bicinchoninic acid assay (BCA) kit (Pierce® BCA assay kit, Thermo Fisher). Protein (30 μg diluted in Laemmli 5× buffer containing 10% b-mercaptheoethanol) was denatured and electrophoresed through 4–20% Mini-PROTEAN® TGX Stain-Free™ gels or 4–15% Criterion™ TGX Stain-Free™ gels (Bio-Rad Laboratories) in running buffer (0.1% SDS, 14.8% glycine in 100 mM Tris-HCl, pH 8.2) at 60 V for 30 min, then 100 V for 60 min. The blot was then transferred onto a PVDF-FL membrane (Millipore, IPFL00010) using a Trans-Blot® Turbo™ Transfer System (Bio-Rad Laboratories) for 10 min at 25 V. Membranes were blocked for 1 hr in 5 % low fat milk powder and 0.2% NaNO_3_ in pH 8.0 Tris-buffered saline with 0.5 % Tween-20 (TBST). Membranes were washed 3x 10 min with TBST then incubated overnight at 4 °C in primary antibodies (rabbit TDP-43, 1:1000, Proteintech, 10782–2-AP) diluted in 3 % BSA (Sigma-Aldrich, A3912) in TBST. The following day membranes were washed 3x 10 min with TBST before being incubated for 1 h with IRDye 680 secondary antibodies (1:10,000, Li-Cor Biosciences) and rhodamine anti-β-actin (1:5000, Bio-Rad, 12004163) in TBST. Blots were washed 3x 10 min with TBST then imaged using the ChemiDoc™ MP (Bio-Rad). Fiji ImageJ was used to analyse the protein concentration by taking the mean grey value of the band of the target protein, subtracting the background fluorescence, then normalising to β-actin for each sample. Graphs presented as fold-change to TDP-43 mouse protein levels.

### Tissue Collection for Histology

10 mo mice were sacrificed via lethal injection (sodium pentobarbitone, 100 mg/kg, IP injection) then transcardially perfused with 0.1 M phosphate buffered saline (PBS). and 4 % PFA in 0.1 M phosphate buffer (PB). Sciatic nerves were dissected between the sciatic notch and where the nerve bifurcates into the tibial and sural nerves. Bilateral sciatic nerves were collected, one to be utilised for immunohistology and the other for electron microscopy (EM). Nerves collected for immunohistology were placed into 4 % PFA in 0.1 M PBS overnight at 4 °C, then washed with 0.1 M PBS x 3 before being transferred into 30 % sucrose in PBS until processing. Tissue was embedded and stored in O.C.T (SciGen) at -80 ° C. Sciatic nerves were serially sectioned (12 μm) longitudinally in sets of 6 slides, 6-12 sections per slide. For EM, sciatic nerve tissue was subsequently fixed in Karnovsky’s buffer (2 % glutaraldehyde and 2 % paraformaldehyde in 0.1 M sodium cacodylate) then washed 3x with 0.1 M sodium cacodylate and stored at 4 °C in 0.1 M sodium cacodylate until processing. Processing occurred in collaboration with the Ramaciotti Centre for Structural Cryo-Electron Microscopy at Monash University who post fixed samples in 2 % OsO4, then transferred samples through a series of concentrated ethanol stages (50 %, 70 %, 90 %, 100 %), to acetone, and finally from diluted (1 resin:3 acetone, 1 resin:1 acetone, 3 resin:1 acetone) to 100 % resin.

### Immunohistochemistry

O.C.T was removed using 0.1 M PBS. Primary antibodies (Caspr, Abcam, Ab34151, 1:200; pan Neurofascin, R&D Systems, AF3235, 1:200; ChAT, Sigma-Aldrich, AMAB91129, 1:250; GFAP, ThermoFisher, 1:200; c-JUN, Cell Signaling Technology, 1:200; SOX10, R&D Systems, 1:200; Iba1 FUJIFILM Wako, 1:300; CD11b, Bio-Rad, 1:100) were incubated for 24-48 h in 10 % NDS in 0.3 % TritonX-PBS. Secondary antibodies (all raised in Donkey and used at 1:200; Jackson Immuno Research; and stains where nuclei were counted, Hoechst, Invitrogen, 1:1000) were incubated for 1.5-2 h at room temperature. Slides were coverslipped using Fluorescent Mounting media (DAKO).

### Image Acquisition & Analysis

All images for immunocytochemistry were acquired using the LSM900 Zeiss Confocal Microscope (Carl Zeiss, Inc.) at 40 x magnification with a PL-APO 40x/1.3 oil immersion lens or 20x magnification with a PL-APO 20x/0.8 air lens. The entire tissue section was captured (z-step size of 1-1.5 μm) and all images were obtained using consistent laser and gain settings. 3-6 images from one slide per mouse were captured, 3-5 mice per genotype were assessed. All images were analysed using Fiji Image J software. Maximum projection images were assessed. Node length was determined using the line tool, with three parallel lines drawn and averaged. Areas of the nodes and paranodes were taken via the polygon selection tool and regions of interest for each gap junction were obtained. The multi-measure tool was used to obtain mean grey values for each paranode and node for protein brightness quantification. Cell counts were obtained using the FIJI Cell Counter Software. For c-jun and GFAP counts, 3 fields of view of 25,000 μm^2^ area each were counted per image, 3x images per animal, 5 animals per genotype. For Iba1 and CD11b counts, images were captured on the microscope using a 20x objective and all positive cells within each field of view were counted, 3x images per animal, 3 animals per genotype.

SCoRe is an imaging technique used to quantify myelin density, due to the unique refractive index of compact myelin (Schain et al., 2014). Three confocal lasers (458 nm, 561 nm, & 633 nm) were utilised to obtain reflection signals and combined to represent whole myelinated axons. All SCoRe images were taken with the LSM780 Zeiss Confocal Microscope (Carl Zeiss, Inc.) at 40x magnification with a PL-APO 40x/1.4 (DIC) oil immersion lens. For analysis, a threshold was selected that best captured bona fide myelin processes yet minimised background noise, and this threshold was maintained for all images. The percentage of positive pixels for the tissue area of interest was analysed using FIJI on 3-12 images per mouse, 5-11 mice per genotype.

### Transmission Electron Microscopy (TEM)

Imaging was performed using the JEOL 1400 flash obtained at 5,000x magnification. 5 images per animal were assessed for both g-ratio and myelin abnormalities (misfolded myelin, myelin non-compaction, and axon or myelin degeneration, as similarly quantified by our lab (Lewis et al., 2025) and others (Kang et al., 2013; Li et al., 2018; Trapp et al., 1982)), 5 animals per genotype were assessed.

### Fluorogold Injections & Histology

7 days prior to sacrifice, male 10 mo TDP-43 (n = 3) and WT (n = 3) mice were administered an IP injection of Fluorogold (2 mg Fluorogold powder (SANTSC-358883) dissolved in 1 mL distilled water). 200-400 μL was administered based on the weight of each mouse. Mice were euthanized via lethal injection (sodium pentobarbitone, 100 mg/kg, IP injection) then transcardially perfused with 0.1 M phosphate buffered saline (PBS). Mice were then transcardially perfused with 4 % PFA in 0.1 M phosphate buffer (PB). Spinal cords were dissected out and placed in 4 % PFA overnight at 4 °C, then washed with 3 x 0.1 M PBS and transferred into 30 % sucrose in PBS. Lumbar spinal cord was dissected and frozen in O.C.T (SciGen). Spinal cords were serially sectioned (25 μm) transversally in sets of 6 slides, 10-12 sections per slide. O.C.T was removed using 0.1 M PBS. Slides were placed in EDTA buffer pH 6 (Tris-HCU pH 8.5 100mM, NaCl 200nM EDTA pH 8.0 5mM, SDS 0.2%, MilliQ) in a 99 °C heat bath for 10 min for antigen retrieval. Slides were washed 3 x 5 min PBS then primary antibodies (ChAT, Sigma-Aldrich, AB144P; Anti-Fluorescent Gold Antibody, Sigma-Aldrich, AB153-I, 1:1000; NeuN, Abcam, ab279296, 1:1000) were incubated overnight at room temperature in 10 % NDS in 0.3 % TritonX-PBS. Secondary antibodies (all raised in Donkey and used at 1:200; Jackson Immuno Research) were incubated for 1.5-2 h at room temperature. Slides were coverslipped using Fluorescent Mounting media (DAKO).

### Sciatic Nerve Electrophysiology

Sciatic nerve electrophysiology was recorded in 10 mo WT (n = 4M; 2F), TDP-43 (n = 4M; 3F), and P0-cre/TDP-43 (n = 4M; 3F) mice. Mice were anaesthetized in 2-3 % isoflurane in 100 % oxygen via nose-cone inhalation on a rodent ventilator (Ugo Basile). Animals were placed prone on a heating pad to maintain body temperature. The left hindquarters were immersed in a bath of liquid paraffin oil (Gold Cross), also on the heating pad, and its temperature was measured with a thermocouple to ensure temperature stayed at 37 °C. The left foot of the mouse was tied and hooked to the bath to keep the leg straight and taut. The sciatic nerve (∼6 mm) and gastrocnemius muscle were exposed, and a small section of plastic sheeting was inserted beneath the sciatic nerve for insulation. Three fine silver wire stimulating electrodes were inserted 2 mm apart through a portion of silastic tubing (diameter 500 μm) with a slit to create a cuff. This cuff was placed underneath the sciatic nerve, ensuring the nerve was in contact with each electrode. A common anodal electrode made of silver wire was inserted proximally above the leg, allowing cathodal stimuli to be delivered to each of the cuff electrodes in turn. A pair of silver wire recording electrodes (separation ∼ 5 mm) was placed on the largest area of exposed gastrocnemius muscle and used to differentially record the muscle action potential (M-wave); a reference silver wire grounding electrode was placed under the skin of the leg. M-wave responses were recorded, amplified, filtered (10–100 Hz high pass, 600–1000 Hz low pass; Neurolog) and digitized at 10 kHz on a computer-based system (CED Power 1401 interface with Spike 2 software; Cambridge Electronic Design). Monophasic square-wave constant voltage pulses (SD9 Square Pulse Stimulator; Grass Instruments) of 0.1 ms duration were delivered at 2 Hz. A series of increasing voltage stimuli was applied to each of the 3 cuff electrodes to measure the voltage giving a maximal m-wave, then further increased by 20 % to ensure supramaximal stimulation (EMTEK 20 MHz Oscilloscope). M-wave responses to 10 supramaximal stimuli to each cuff electrode were averaged and used for analysis. For CMAP amplitude, onset latency, area under the curve, and duration measurements, response to the 3 electrodes were combined and averaged per animal. All data were analysed according to established electrophysiology practice (Barkhaus et al., 2024; Kouzaki et al., 2016; Mallik & Weir, 2005). For conduction velocity measurements, M-wave responses to each electrode were examined individually, whereby the half maximal point of the rising phase of the M-wave was recorded. Since there was a known distance of 2 mm between each electrode, conduction velocity was calculated using the time delay between each electrode’s M-wave for each animal. After recordings were obtained, animals remained anesthetized and were sacrificed via lethal injection. All data was exported from Spike2 to Microsoft Excel and analysed in GraphPad Prism.

### *In vivo* Axon Transport Imaging

Axonal transport assessments were performed in 10 mo WT (n = 3M; 3F), TDP-43 (n = 3M; 3F), and P0-cre/TDP-43 (n = 3M; 3F) and *in vivo* signalling endosomes were captured using methodology previously described (Sleigh, Tosolini, & Schiavo, 2020; Tosolini et al., 2021) using the AlexaFluor 555 fluorescently tagged tetanus neurotoxin (HcT-555) that was a generous gift from Giampietro Schiavo and James Sleigh (University College London, UK). Briefly, HcT (residues 875–1315) were fused to an improved cysteine-rich region was bacterially expressed as a glutathione-S-transferase fusion protein (Restani et al., 2012), then cleaved and labelled with AlexaFluor555 C_2_ maleimide (Thermo Fisher Scientific, A-20346). Mice were anaesthetised with 2-3 % isoflurane in 100% oxygen using a rodent ventilator (Ugo Basile). On the left leg, a 1-2 mm long skin incision was made over the tibialis anterior (TA) muscle. 3-5 μL of HcT-555 in PBS (7.5–10 μg of H_C_T-555 in PBS) was injected into the TA muscle using pulled graduated, glass micropipettes (Drummond Scientific,5-000-1001-X10) (Rahul Mohan et al., 2015) then the skin was sutured, and mice were left to recover. 4 h post injection, mice were again anaesthetised (2-3 % isoflurane in 100 % oxygen) and the sciatic nerve of the injected (left) leg was exposed. A small piece of parafilm was inserted to separate the sciatic nerve from the underlying connective tissue/musculature, before the mouse and anaesthetic nosepiece was transferred to a customized stage on an inverted LSM900 confocal microscope (Carl Zeiss Inc.) enclosed within an environmental chamber maintained in a 37 °C chamber. A 40x Plan-Apochromat oil immersion objective lens (Zeiss) was utilised allowing the imaging of an area of the nerve containing axons transporting the HcT-555. Videos were obtained between 1-1.25 frames per second and were taken on a single plane of focus per video at <1 % laser power. Animals were ethically sacrificed after imaging via cervical dislocation. Analysis of signalling endosomes was performed using Fiji Image J via the TrackMate plugin (Kinetic Imaging Ltd) (Ershov et al., 2022). A minimum of 20 retrogradely travelling signalling endosomes that could be viewed for > 15 consecutive frames and did not exhibit > 10 consecutive pauses were included for analysis. A pause was considered a signalling endosome that remained stationary (defined as a speed of less than 0.01 m/s). A minimum of 3 axons per animal were assessed, thus > 60 signalling endosomes per animal.

### Statistical Analyses

All data are expressed as the mean ± the standard error of the mean (SEM) unless stated otherwise. The sample sizes (*n*) are specified in each figure. Multiple comparisons corrections were performed when appropriate and stated in each figure legend. All data were analysed using GraphPad Prism 9 or 10 software. A statistically significant difference was concluded if p < 0.05. Statistical significance intervals were identified as follows: *p < 0.05, **p < 0.01, ***p < 0.001, ****p < 0.0001. P values between 0.05 and 0.15 were considered trending. For ANOVAs and Mixed-Effects models, p-values shown are the post-hoc tests only. If post-hoc p-values are present, ANOVA p-values were statistically significant (*p < 0.05).

## Author Contributions

Designed research: KNL, SKB, BJT & DGG. Performed research: KNL, AA, GC, APT, RM. Analyzed data: KNL. Wrote the first draft of the paper: KNL. Wrote the paper: KNL, SKB. Edited the paper: KNL, AA, GC, APT, RM, STN, DGG, BJT, SKB.

## Acknowledgments

The authors acknowledge the Florey Neuroscience Microscopy Facility, the Integrated Physiology Facility (SBMS, UQ), and the Microscopy and Image Analysis Facility (SBMS, UQ). We acknowledge the Monash Centre for Electron Microscopy for the TEM processing. This work was supported by an MND Research Australia Superball XVI MND Research Grant (IG 2403). S.B. was supported by a Rebecca L. Cooper Al & Val Rosenstrauss Medical Research Fellowship. B.T. was supported by FightMND Drug Screening Program and Stafford Fox Medical Research Foundation Grants. K.L. was supported by an Australian Government Research Training Scholarship and Stipend and MND Research Australia PhD Top-Up Scholarship. A.P.T was supported by Junior Non-Clinical Project Grant from the Motor Neuron Disease Association (Tosolini/Oct20/973799); an MND Research Australia Col Bambrick MND Research Grant (IG 2450). S.T.N was supported by a FightMND Drug Development Grant (DDG-73) and the Australian Institute for Bioengineering and Nanotechnology.

---

**Supplementary Figure 1.**
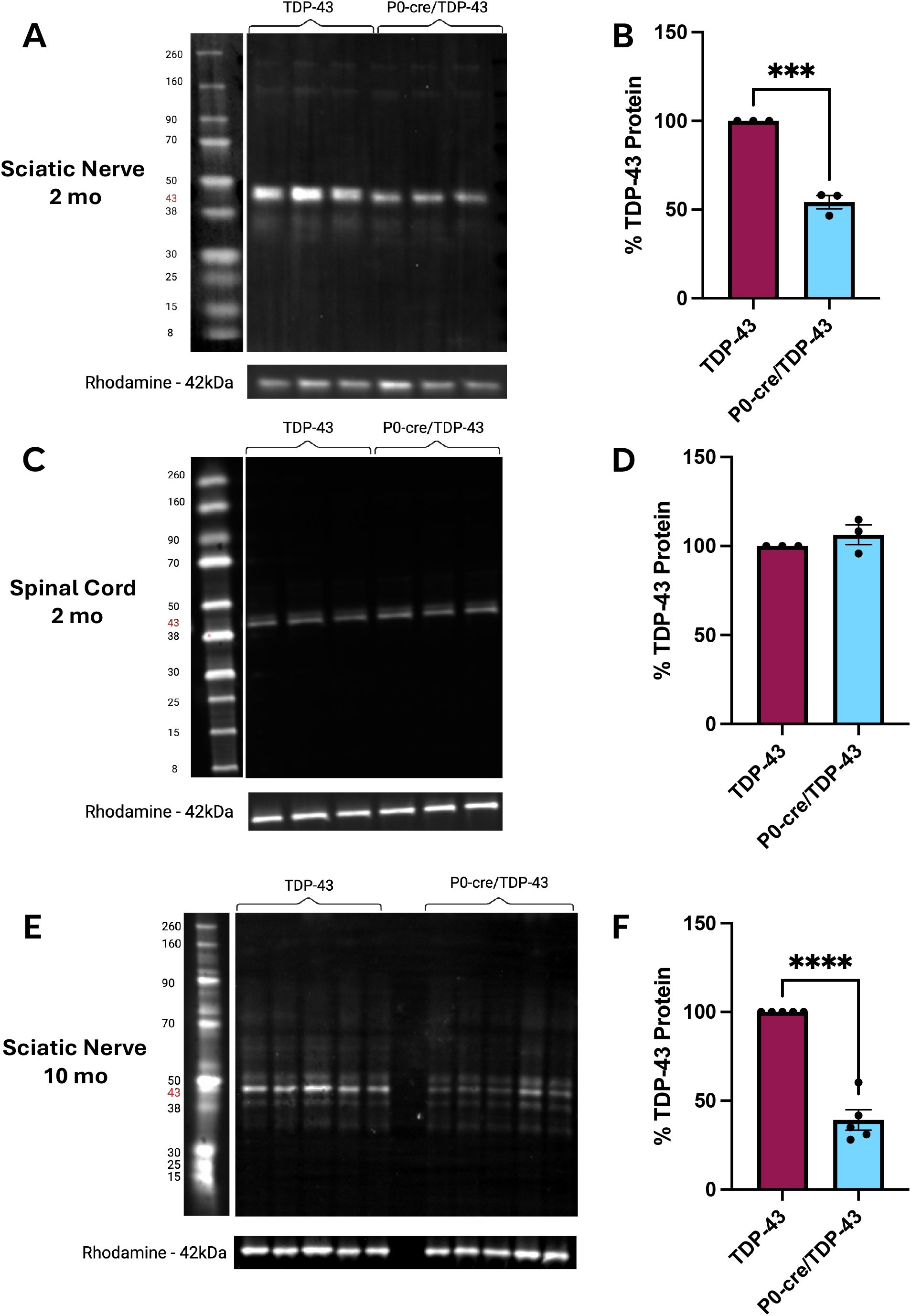
TDP-43 protein analysis in 2 mo and 10 mo TDP-43 and P0-cre/TDP-43 mice. **A.** Western blotting of sciatic nerves revealed TDP-43 protein was significantly reduced in 2 mo P0-cre/TDP-43 mice compared to TDP-43 mice (**B**), with no changes in the spinal cord (**C, D**). Western blotting of sciatic nerve TDP-43 protein remained depleted in 10 mo TDP-43 mice compared to P0-cre/TDP-43 (**E, F**). Mean TDP-43 Protein Fold Change ± S.E.M, *p<0.05, ***p<0.001, ****p<0.0001, Student’s unpaired t-tests, n = 3-5 mice per genotype.

**Supplementary Figure 2.**
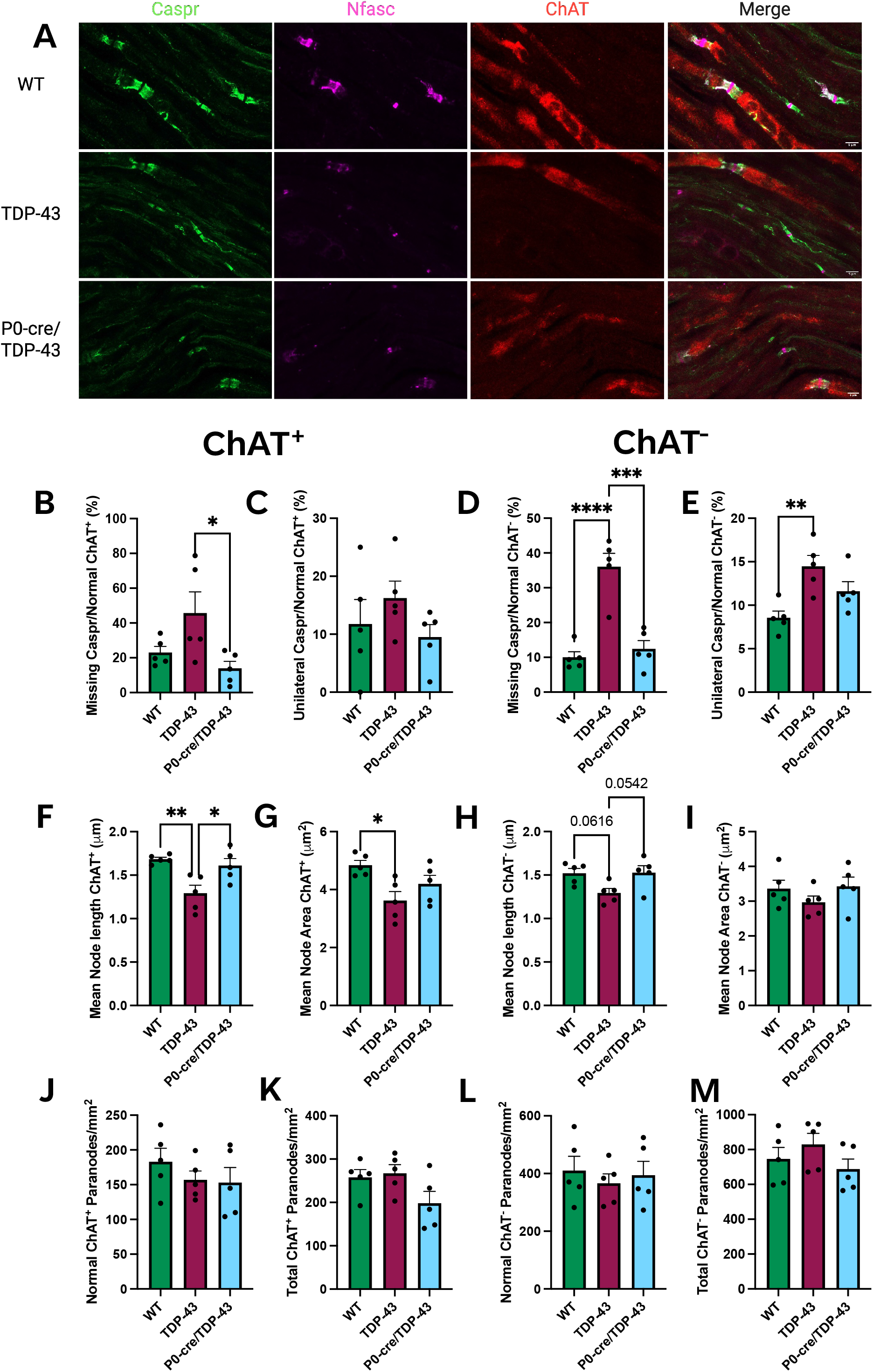
Node and paranode structure and size comparisons between motor and sensory neurons in sciatic nerves of 10 mo mice. **A.** Representative images of the nodes and paranodes of WT, TDP-43, and P0-cre/TDP-43 mice sciatic nerves, with contactin-associated protein (Caspr; green) marking the paranodes, Neurofascin (Nfasc; magenta) at the node and some paranodes, and choline acetyltransferase (ChAT; red) marking motor neurons, scale bars 5 µm. **B.** P0-cre/TDP-43 mice had significantly less missing Caspr^+^ paranodes compared to TDP-43 (p = 0.0314), while for ChAT^-^ neurons, TDP-43 mice exhibited a significant increase in missing paranodes compared to WT (**D**; p < 0.0001) and P0-cre/TDP-43 (p = 0.0002). **C.** The ratio of unilateral ChAT^+^ paranodes to normal was unchanged between genotypes, however this ratio for ChAT^-^ paranodes was significantly increased for TDP-43 compared to WT (**E**; p = 0.0048) **F.** Mean node length for ChAT^+^ nodes was significantly decreased in TDP-43 mice compared to WT (p = 0.0060) and was rescued for P0-cre/TDP-43 (p = 0.0217), while for ChAT^-^ neurons, there was a trend of decreased node length for TDP-43 compared to WT (**H**; p = 0.0616) and a trend of a rescue between TDP-43 and P0-cre/TDP-43 (p = 0.0542). **G.** Mean node area was significantly decreased in ChAT^+^ neurons for TDP-43 compared to WT (p = 0.0180) but was no different between genotypes for ChAT^-^ neurons (**I**). The density of normal paranodes for both ChAT^+^ (**J**) and ChAT^-^ (**L**) axons was unchanged between genotypes. The density of total paranodes was unchanged between genotypes for ChAT^+^ axons (**K**) and ChAT^-^axons (**M**). All data are presented as mean ± S.E.M, *p < 0.05, **p < 0.01; ***p < 0.001; ****p < 0.0001; One-way ANOVA with Tukey’s *post hoc* test; n = 5 mice per genotype, 100-200 nodes and paranodes analysed per animal.

**Supplementary Figure 3.**
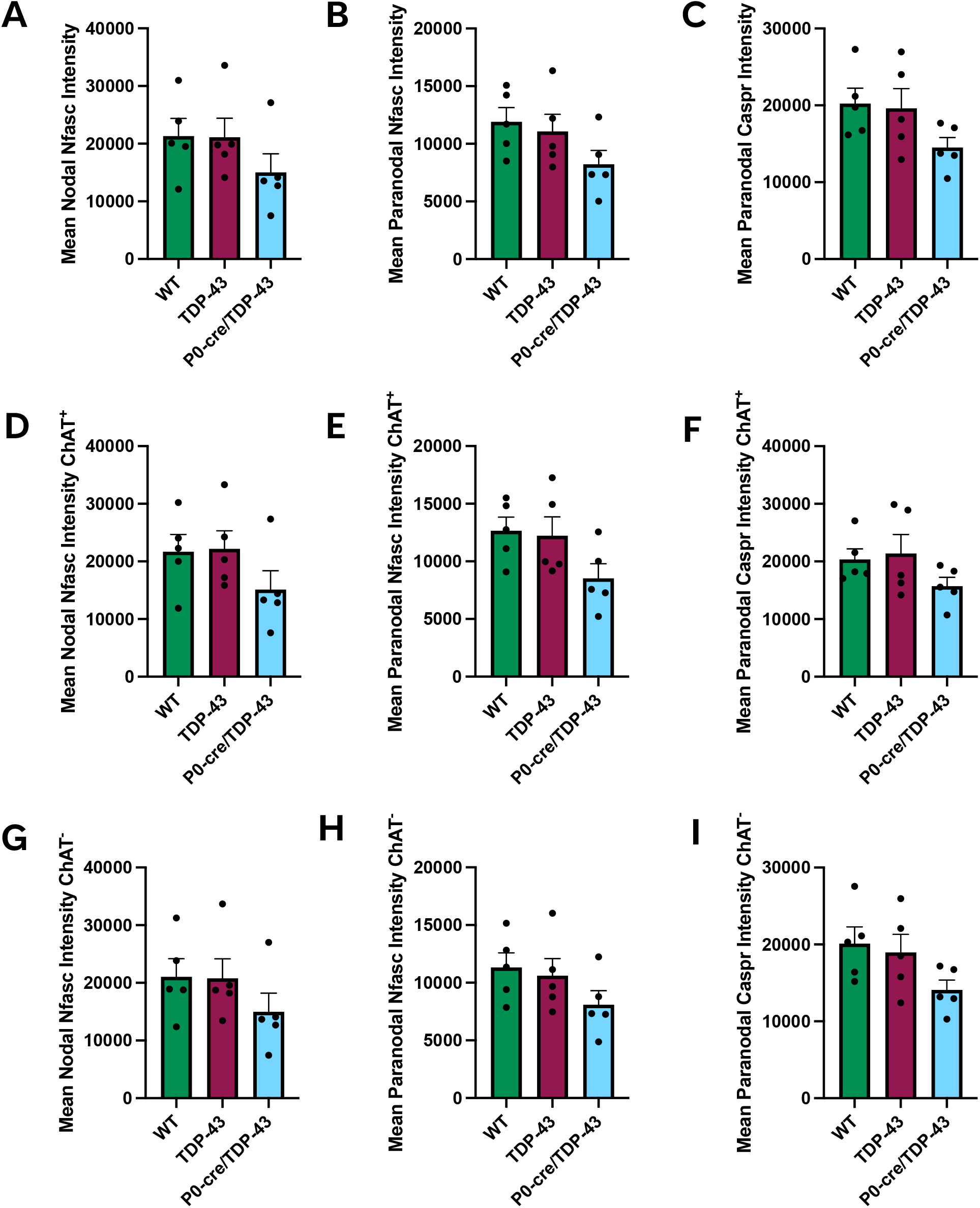
Protein staining intensity at the nodes and paranodes in the sciatic nerves of 10 mo mice. Neurofascin (Nfasc) was measured at the node and paranodes, while contactin-associated protein (Caspr) was measured at the paranodes. There were no changes between genotypes in the mean intensity of Nfasc staining at the nodes (**A**), nor when stratified for ChAT^+^ (**D**) and ChAT^-^ (**G**) neurons. Mean paranodal Nfasc staining intensity was unchanged between genotypes (**B**) however in ChAT^+^ paranodes, there was a trend for decreased intensity for P0-cre/TDP-43 mice compared to WT (**E**; p = 0.1248), while there was no change between genotypes for ChAT^-^ paranodes (**H**). Mean paranode Caspr intensity was unchanged between genotypes (**C**) and for ChAT^+^ neurons (**F**), with a decreased trend for P0-cre/TDP-43 compared to WT at ChAT^-^ neurons (**I**; p = 0.1237). All data are presented as mean ± S.E.M, *p < 0.05; One-way ANOVA with Tukey’s post hoc test; n = 5 mice per genotype, 100-200 nodes and paranodes analysed per animal.

**Supplementary Figure 4.**
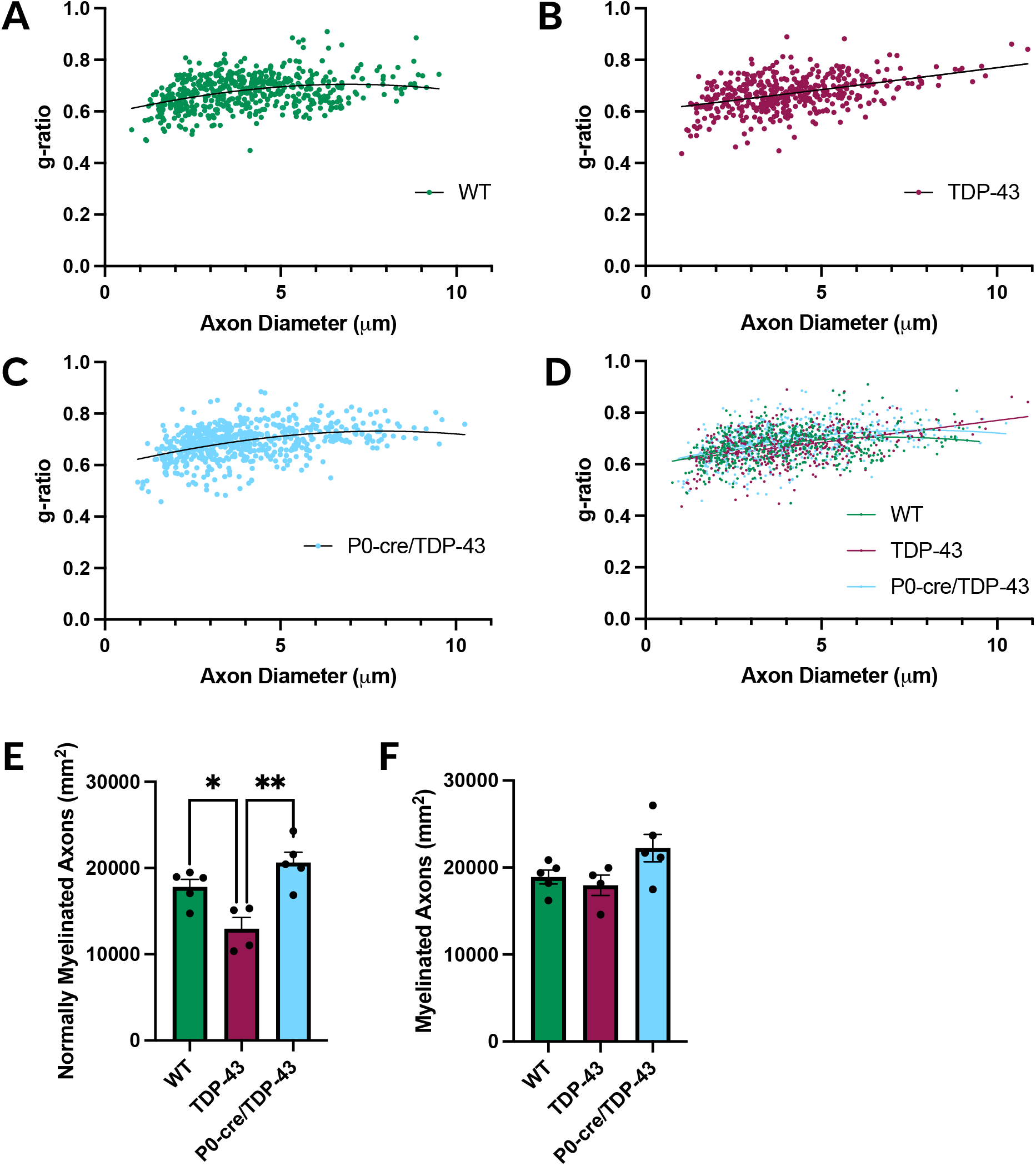
The Densities of Myelinated Axons and G-ratio Spreads of WT, TDP-43, and P0-cre/TDP-43 Mice. Scatter plots of g-ratio vs. axon diameter for WT (**C**), TDP-43 (**D**), P0-cre/TDP-43 (**E**), and combined (**F**) showed no significant differences in the spread of values. **A.** The density of normally myelinated axons in 10 mo sciatic nerves was significantly decreased in TDP-43 compared to WT (p = 0.0302) and P0-cre/TDP-43 (p = 0.0017). **B.** The density of all myelinated axons (combination of normally and abnormally myelinated axons) was unchanged between the three genotypes at 10 mo. Data are presented as mean ± S.E.M or scatter plots, *p < 0.05, ** p < 0.01; One-way ANOVA with Tukey’s post hoc test; n = 4-5 mice per genotype, 100-150 axons analysed per animal.

**Supplementary Figure 5.**
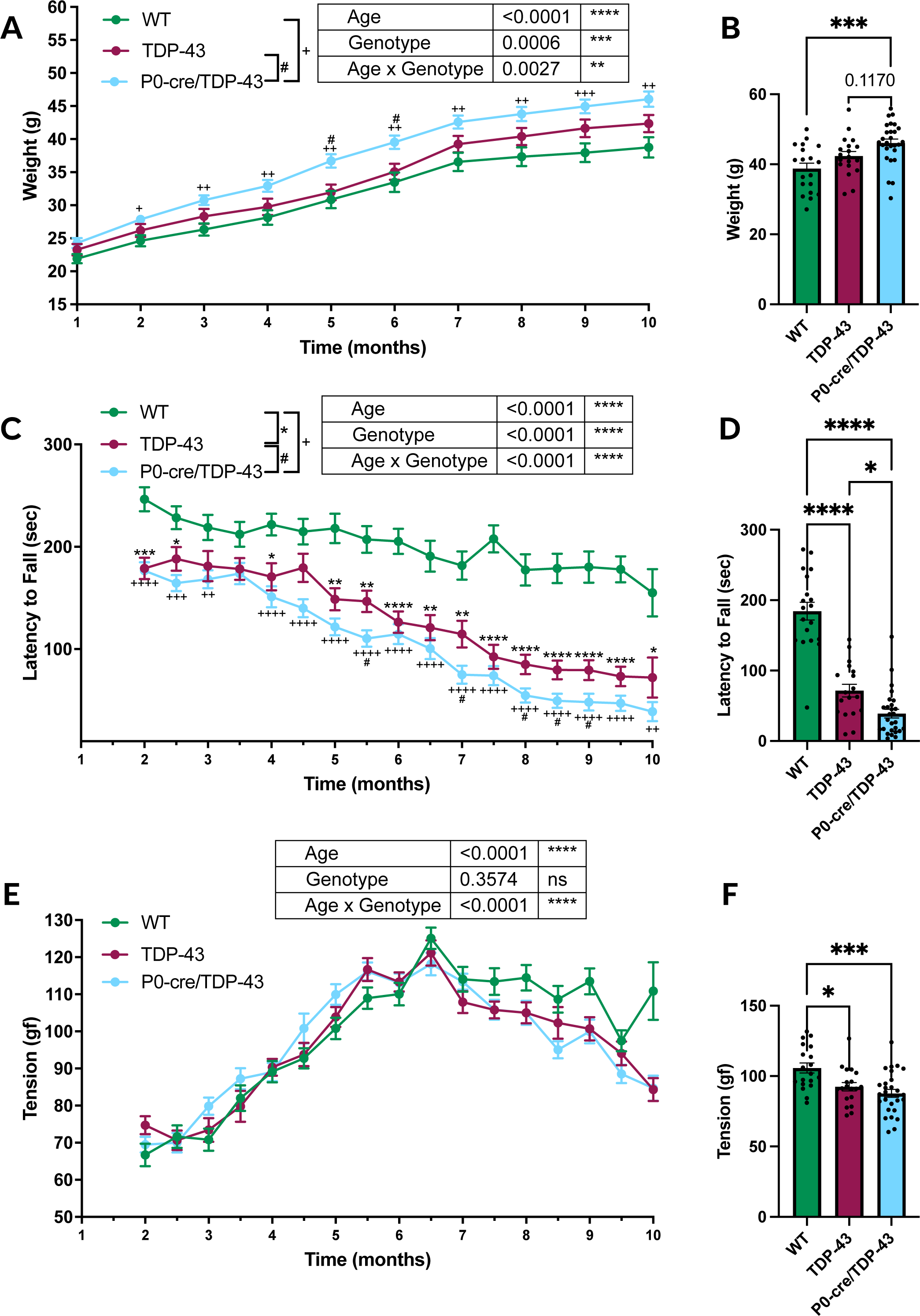
Motor performance, including hind-limb grip strength and the rotarod walking test, was tested throughout the disease course from 2 mo to 10 mo in WT, TDP-43, and P0-cre/TDP-43 mice. **A.** P0-cre/TDP-43 mice weighed significantly more than WT (p = 0.0006), with P0-cre/TDP-43 mice being significantly heavier than WT throughout the disease course (2-10 mo) and significantly heavier than TDP-43 at 5 and 6 mo. **B.** At 10 mo, P0-cre/TDP-43 mice were significantly heavier than WT (p = 0.0005), with a trend towards increased compared to TDP-43 (p = 0.1170). **C**. The rotarod walking test revealed that TDP-43 and P0-cre/TDP-43 mice fell faster than WT (p < 0.0001). TDP-43 mice fell faster than WT at 2 and 2.5, then from 4-10 mo, while P0-cre/TDP-43 mice fell faster than WT from 2-3 mo, then 4-10 mo, and fell faster than TDP-43 at 5.5, 7, 8, 8.5, and 9 mo. **D.** At 10 mo TDP-43 and P0-cre/TDP-43 fell faster than WT (p < 0.0001 and p < 0.0001, respectively) and P0-cre/TDP-43 fell faster than TDP-43 (p = 0.0335). **E.** Hind-limb grip strength analysis revealed no longitudinal changes between genotype across disease course, but at 10 mo a significant reduction in grip strength for TDP-43 and P0-cre/TDP-43 compared to WT (**F**; p = 0.0172 and p = 0.0003, respectively). All data are presented as mean ± S.E.M, Mixed effects analysis with Tukey’s post-hoc analyses for line graphs, One-Way ANOVAs for 10 mo analyses, n = 19-28 (12 F; 8-16 M) per genotype, *p<0.05, **p<0.01, ***p<0.001, ****p<0.0001.

**Supplementary Figure 6.**
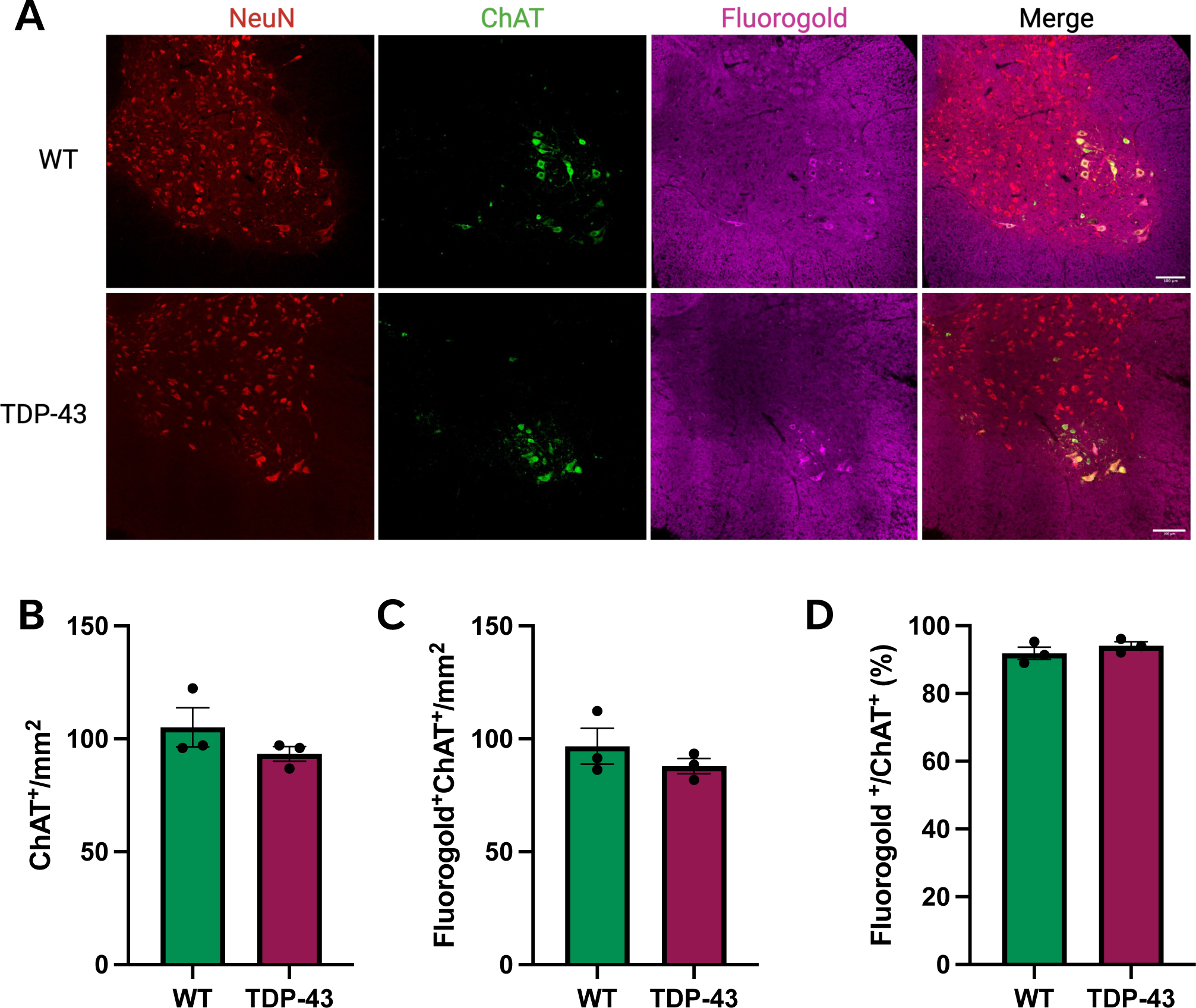
Assessment of lumbar spinal cord motor neurons using retrograde tracer fluorogold, administered 7 days prior to collection in WT and TDP-43 10 mo mice. **A.** Representative images of the ventral horn grey matter of lumbar spinal cord cross sections for both WT and TDP-43. All neurons are marked with NeuN (red), motor neurons are labelled with choline acetyltransferase (ChAT; green), and motor neurons with intact axons with fluorogold (magenta), scale bars 100 µm. Between WT and TDP-43, were no differences in the density of ChAT^+^ motor neurons (**B**), co-labelled ChAT^+^ motor neurons and fluorogold^+^ motor neurons (**C**), nor the ratio of fluorogold^+^ motor neurons to ChAT^+^ motor neurons (**D**). All data are presented as mean ± S.E.M, student’s unpaired t-tests, n = 3 per genotype, *p<0.05.

**Supplementary Figure 7.**
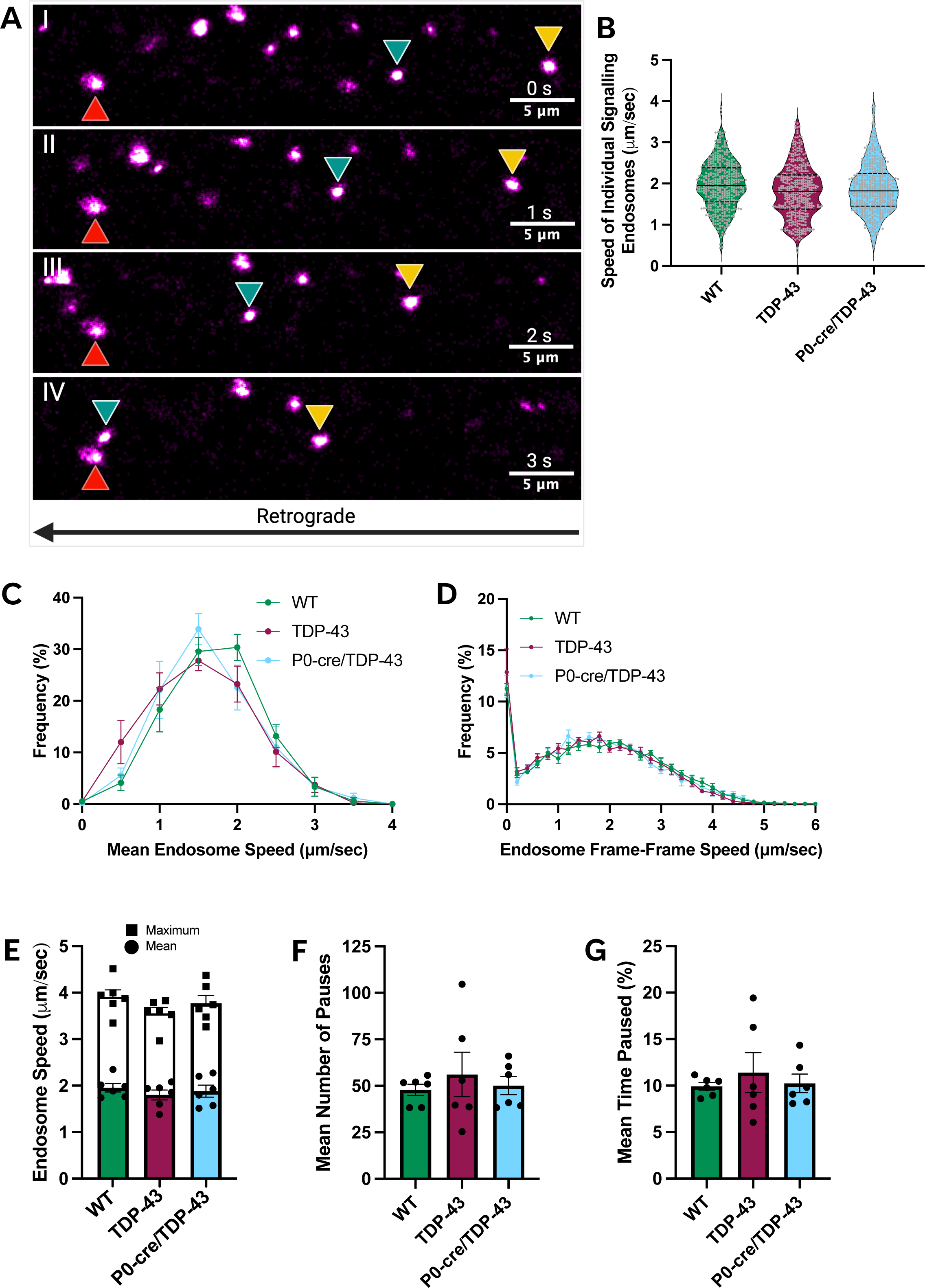
*In vivo* imaging of retrograde signalling endosome axonal in 10 mo WT, TDP-43, and P0-cre/TDP-43 mice. **A.** Representative images of signaling endosomes labelled with tetanus toxin fragment (HcT-555; magenta). Arrowheads depict signaling endosomes that are moving retrogradely (i.e. right to left; panel i at time zero, ii at 1 s, iii at 2 s, and iii at 3 s) along the neuronal axon (yellow and blue arrow) or endosomes that are stationary or paused (red). Videos were acquired at 1-1.25 frames per second. Individual signalling endosome speeds (**B**), the frequency distribution of mean endosome speeds (**C**), and frame-toframe endosome speeds (**D**) were comparable across genotypes. Similarly, no differences were observed in the mean or maximum signaling endosome speeds (**E**), mean number of pauses (**F**), or time paused (**G**) between WT, TDP-43, and P0-cre/TDP-43 mice. One-Way ANOVAs, *p < 0.05; n = 6 mice (3 F; 3 M) per genotype, 3 axons per mouse, >20 endosomes tracked per axon.

